# A Unified Catalog of 19,251 Non-human Reference Species Genomes Provides New Insights into the Mammalian Gut Microbiomes

**DOI:** 10.1101/2022.05.16.491731

**Authors:** Xiaoping Li, Chen Tian, Daohua Zhuang, Liu Tian, Xingwei Shi, Yanli Bai, Han Gao, Hong Zhou, Fangfang Zhao, Min Dai, Lei Zhu, Qunfu Wu, Xiaotong Liu, Tao Zhang, Jianan Sang, Sunil Kumar Sahu, Xun Xu, Huijue Jia, Huan Liu, Liang Xiao, Karsten Kristiansen, Zhigang Zhang

**Affiliations:** State Key Laboratory for Conservation and Utilization of Bio-Resources in Yunnan, School of Life Sciences, Yunnan University, Kunming, Yunnan 650091, P.R. China; BGI-Shenzhen, Shenzhen, Guangdong 518083, P.R. China; Laboratory of Genomics and Molecular Biomedicine, Department of Biology, University of Copenhagen, Copenhagen 2100, Denmark; Institute of Metagenomics, Qingdao-Europe Advanced Institute for Life Sciences, BGI-Qingdao, Qingdao, Shangdong166555, P. R. China

## Abstract

The gut microbiota is essential for host health and survival. Here, using samples from animals living in the Qinghai-Tibetan Plateau, we recovered 119,568 metagenome-assembled genomes (MAGs) that were clustered into 19,251 species-level genome bins (SGBs) of which most represent novel species. We present a novel mechanism shaping mammalian gut microbiomes using ancestral founder bacteria (AFB) as a core skeleton and recurring lineage-specific gains of microbial species that are transferred frequently among multiple hosts, not strictly limited by host phylogeny. Such lineage specific gains are responsible for increasing gut microbial diversity, maintaining functional stability, and endowing specific functions for host adaptions. Our analyses did not support the existence of co-phylogeny or co-speciation events between mammal hosts and their individual gut symbionts. The results presented in this study not only reveal novel unique gut microbial species and offer insight of value for understanding the diversity, stability, functionality of the mammalian gut microbiomes, and the co-evolution with their hosts, but also emphasize that animals living in extreme environments are a promising resource for the discovery of novel biological functions.

---

The gut microbiota constitutes an essential functional unit of the mammalian body involved in nutrient utilization, immune development, and host survival in extreme environments *(1-5)*. However, except for the gut microbiota of humans *(6, 7)*, the diversity, stability, and functional traits of mammalian gut microbiomes and the co-evolution with their hosts are poorly understood in part due to lack of comprehensive microbial reference genomes. Recently, 1,209 species-level genome bins (SGBs) were identified by analysis of the gut microbiota from 184 unique species representing five main groups of animals *(8)*. Of these SGBs, 75% represented novel microbial species testifying to the still huge uncharted domains of the gut microbiota. Thus, in-depth investigation of the gut microbiota of diverse non-human mammals, including animals living in extreme environments such as mammals living on the Qinghai-Tibetan Plateau is likely to uncover novel gut microbial species. The Qinghai-Tibetan Plateau, also termed “the third pole”*(9)*, represents a natural laboratory for studying evolution and environmental changes *(10)* and can be considered as an evolutionary junction for the history of modern biodiversity *(11)* and ice age megaherbivores*(12)*.

Numerous studies have demonstrated the influence of host genetics on the gut microbiome *(13)*, and vertical maternal-to-offspring transmission is key to maintain host-population-level stability of the gut microbiota *(14-16)*. Further observations suggest that microbial community relationships parallel the phylogeny of their hosts, coined phylosymbiosis *(17)*. Phylosymbiosis represents a simple ecological modeling of host filtering *(18)*, with consequences that we can observe now, but phylosymbiosis is unable to explain the long-term co-evolutionary mechanisms and dynamics of the gut microbiota and their mammalian hosts *(19)*. Few phylogenetic analyses have reported that co-speciation plays a predominant role in the co-evolution between mammals and their gut commensals, but the conclusions are tenuous, because frequent horizontal gene transfer (HGT) events across prokaryotes cause an inaccurate estimation of their phylogenies that cannot be conquered by the limited phylogenetic information provided by analysis of partial 16S rRNA gene sequences or single-copy genes *(20-24)*. Accordingly, whole microbial genome information is needed to improve analyses *(25)* and will be required to unravel the evolutionary dynamics of the gut microbiome’s function during host evolution.

Here, we report results from metagenomic deep sequencing of fecal samples obtained from 1,412 individuals of six high-altitude herbivorous mammals belonging to the two sister orders Perissodactyla and Artiodactyla freely living on the Qinghai-Tibet Plateau.

## Results

### Recovering 119,568 microbial genomes from six non-human mammals

A total of 1,412 fresh fecal samples were collected from six non-human mammals including Yak (388), Tibetan antelope (255), Tibetan cattle (196), Tibetan sheep (446), Tibetan horse (79) and Tibetan Ass (48) from the Qinghai-Tibet plateau (Fig. 1A and Supplementary Table 1). The animal hosts have a divergence time of ∼ 78 Mya (million years ago) representing the divergence of the Perissodactyla and the Artiodactyla orders based on genome inferences *(26-29)*. The integrated pipeline for constructing the bacterial genomes and gene catalogs is shown in Fig. 1B. After DNA extraction and whole genome sequencing, more than 2.23×10^11^ 150 bp paired end reads were produced corresponding to a total of 33.55 Tb raw data (23.74±7.22Gb per sample) (Supplementary Text; Supplementary Table 2). To maximize assembly of MAGs, we employed a co-binning strategy to reconstruct bacterial and archaeal genomes from microbial communities according to tetranucleotide frequency, and abundance correlations of contigs in multiple samples (Supplementary Methods). We evaluated this strategy on the 79 Tibetan horse samples with different multiple sample size settings (Supplementary Text; Supplementary Fig. 1). To standardize the genome quality across other sets, MAGs were retained with >50% genome completeness and <10% contamination, combined with an estimated quality score (QS) (= completeness -5 × contamination) > 50*(30)*. Finally, we recalled a total of 119,568 MAGs from the six animal species, representing 39,278 MAGs from Sheep, 28,125 MAGs from Yak, 10,630 MAGs from Cattle, 8,144 MAGs from Horse, 6,684 MAGs from Ass, and 26,607 MAGs from Tibetan antelope. On average more than 90 MAGs were recovered per sample except for cattle (54) and yak (72). Of these MAGs, 34,977 (29.25%) matched the high-quality genome criterion of >90% completeness and <5% contamination (Supplementary Table 3 and Supplementary Text).

**Figure 1.**
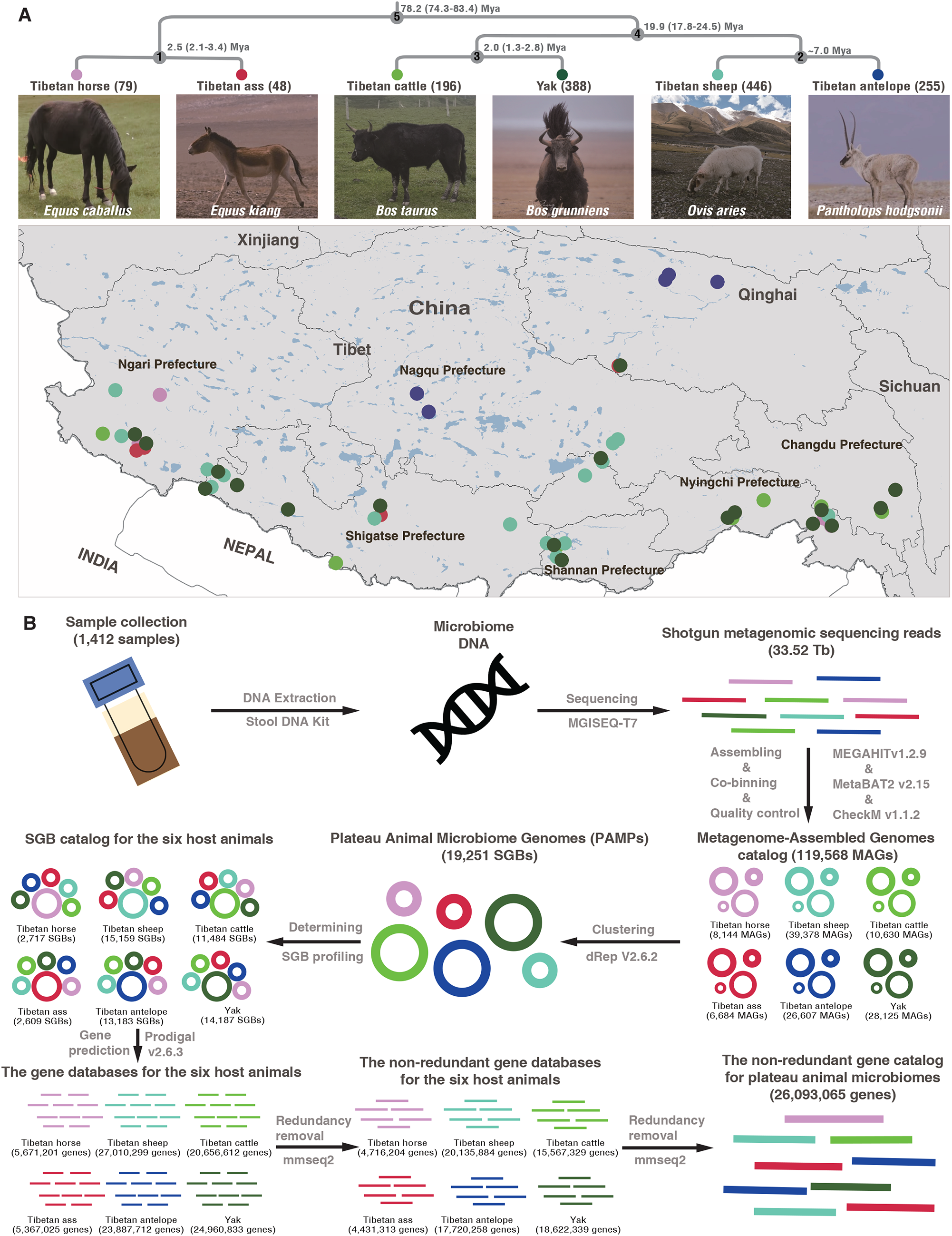
Sample collection and the pipeline for data retrieval. (**A**) The phylogenetic tree of six host animal species living in the Qinghai-Tibet Plateau and geographical distribution of collected fecal samples. (**B**) The pipeline for constructing the non-redundant genome catalog and gene catalog of the non-human mammalian gut microbiomes.

### Reconstructing a catalog of species-level microbial reference genomes from non-human mammalians

According to a threshold of ≥ 95% ANI *(30)* (Supplementary Methods), we clustered all the MAGs obtaining a total of 19,251 SGBs (Supplementary Table 4). Among these, 7,652 (39.75%) met the criterion for a high-quality genome with the genome sizes ranging from 0.53Mb to 6.14Mb (Supplementary Fig. 2, A to C). About 60.13% of SGBs were supported by at least two conspecific genomes (non-singleton SGBs) (Supplementary Fig. 2D). Rarefaction analysis (Fig. 2A) indicated that the non-singleton species were close to saturation, suggesting that most common microbes in the samples were recalled. We performed SGBs profiling of all 1,412 samples (Supplementary Table 5) based on the relative abundances of species genomes in each sample (Supplementary Methods). A non-redundant gene catalog (26,093,065) was constructed based on the predicted 34,469,579 full length genes (Supplementary Methods, Supplementary Table 6).

**Figure 2.**
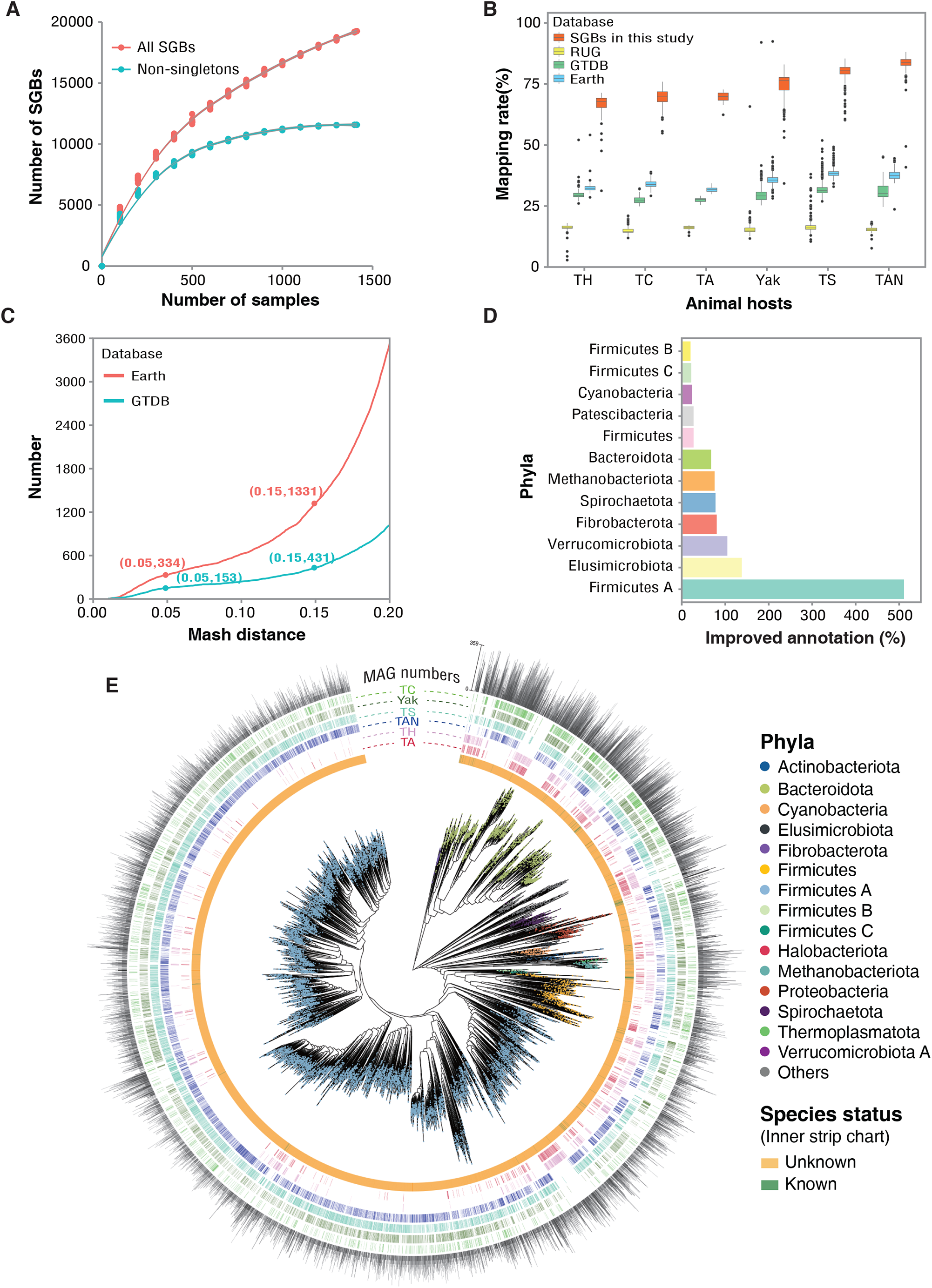
The characteristics and phylogenetic tree of our non-human mammalians microbial reference genomes dataset. (**A**) Rarefaction curve depicting the species diversity assessment of 19,251 SGBs obtained from all 1,412 samples. The curves for all SGBs (red line) and the non-singleton SGBs (blue line) were created by randomly re-sampling the pool of 1,412 samples 10 times with 100 sampling intervals. (**B**) The mapping rate of metagenomic reads from the six host animals against our genome database and other three public reference databases. (**C**) The distribution of mash distances among our identified species and other known species released by the Genome Taxonomy database (GTDB 05-RS95) and Earth Microbiome Project *(33)*. The number of SGB for which mash distance is less than 0.05 and 0.15 is listed in the brackets. (**D**) The expansion of microbial species diversity at the phylum level using the GTDB database *(31)* as the reference. (**E**) The phylogenetic tree of our SGBs was built based on alignment of more than 80 microbial maker genes using the PhyloPhlAn software (See Supplementary Methods) using the default parameters. The 18,607 SGBs containing more than 80 marker genes are shown in the tree. The color of the branches indicates the classification information (See the detailed classification information in Supplementary Table 7). The inner strip shows whether an SGB is unknown (novel identified) or known. The next six strip charts display those SGBs appearing in each of the six host animal gut microbiomes. The outer strip chart shows the MAGs number that supports the SGBs.

The Genome Taxonomy Database (GTDB release95)*(31)* was used to perform taxonomic annotations (Supplementary Methods). In total, 19,068 bacterial SGBs were identified, but only 142 (0.74%) SGBs were classified as known species, suggesting that our database represented a large number of unknown microbial species. Firmicute A (70.85%) and Bacteroidota (13.59%) were the dominant taxa of our SGBs (Supplementary Fig. 3A). We assembled 183 archaea genomes, all being clearly assigned to three phyla, Methanobacteriota, Thermoplasmatota and Halobacteriota (Supplementary Fig. 3B) and of these archaea genomes 172 (94%) represented novel archaea species.

The average mapping rates of our samples to three databases including the GTDB *(31)*, the Hungate collection *(32)*, and Earth’s Microbiomes catalog (GEM)*(33)* were only 15.80%, 30.23% and 36.44%, respectively (Fig. 2B) (Supplementary Methods). These results pointed a remarkable large number of unknown species, also supported by the mash distance analysis (Fig. 2C). Using our database, the average mapping rate substantially increased by nearly 2-fold, reaching 76.58%, indicating that the sequencing depth of our samples was sufficient to cover the majority of the microbial diversity in our samples.

Many clades including some common phyla in the gut microbiome were largely expanded by our catalog. Compared with the most representative databases GTDB*(31)*, 19,098 species (99.10% bacteria) were identified as novel species. The Firmicutes A, the dominant gut bacterial phylum, includes 2,636 species in the GTDB, compared to 13,509 assembled SGBs belonging to this phylum in our database. Except for 34 known bacterial species, our catalog harbors 13,475 novel species representing a more than 500% increase in annotation alone in this phylum. Interestingly, the order Oscillospirales, reported to be enriched in the herbivore gut microbiome *(34, 35)*, contributed with 8,408 species reference genomes in our catalog, representing a 10-fold increasing compare to the reference genomes of this order in the GTDB. For two rare phyla, Elusimicrobiota and Verrucomicrobiota, the assembled SGBs also added a significant number of species in these two phyla (Fig. 2D and Supplementary Table 7). Finally, 18,607 SGBs covering more than 80 marker genes are displayed in the phylogenetic tree (Fig. 2E).

### Features and evolutionary dynamics of the gut microbiomes in the six host species

The results (Supplementary Text; Fig. 3A-3C; Supplementary Fig. 4) from both alpha-diversity and beta-diversity measures consistently demonstrated that microbial communities largely recapitulate host phylogeny (Fig. 3D; Supplementary Fig. 5), which is compatible with the “Phylosymbiosis” hypothesis *(17, 20)*. To further investigate how the gut microbial species co-evolve with their hosts, we performed an ancestral microbiome reconstruction based on the core microbiome of the six animal hosts using the asymmetrical Wagner parsimony approach in the Count software (v.10.04)*(36)* as used previously *(37)*. Our results clearly demonstrated the evolutionary dynamics of gut microbiomes along the host phylogeny, including the appearance of “ancestral” founder bacteria (AFB), the gain or loss of host-specific gained (HSG) SGBs, and host shared (HS) SGBs among the host species (Fig. 4A).

**Figure 3.**
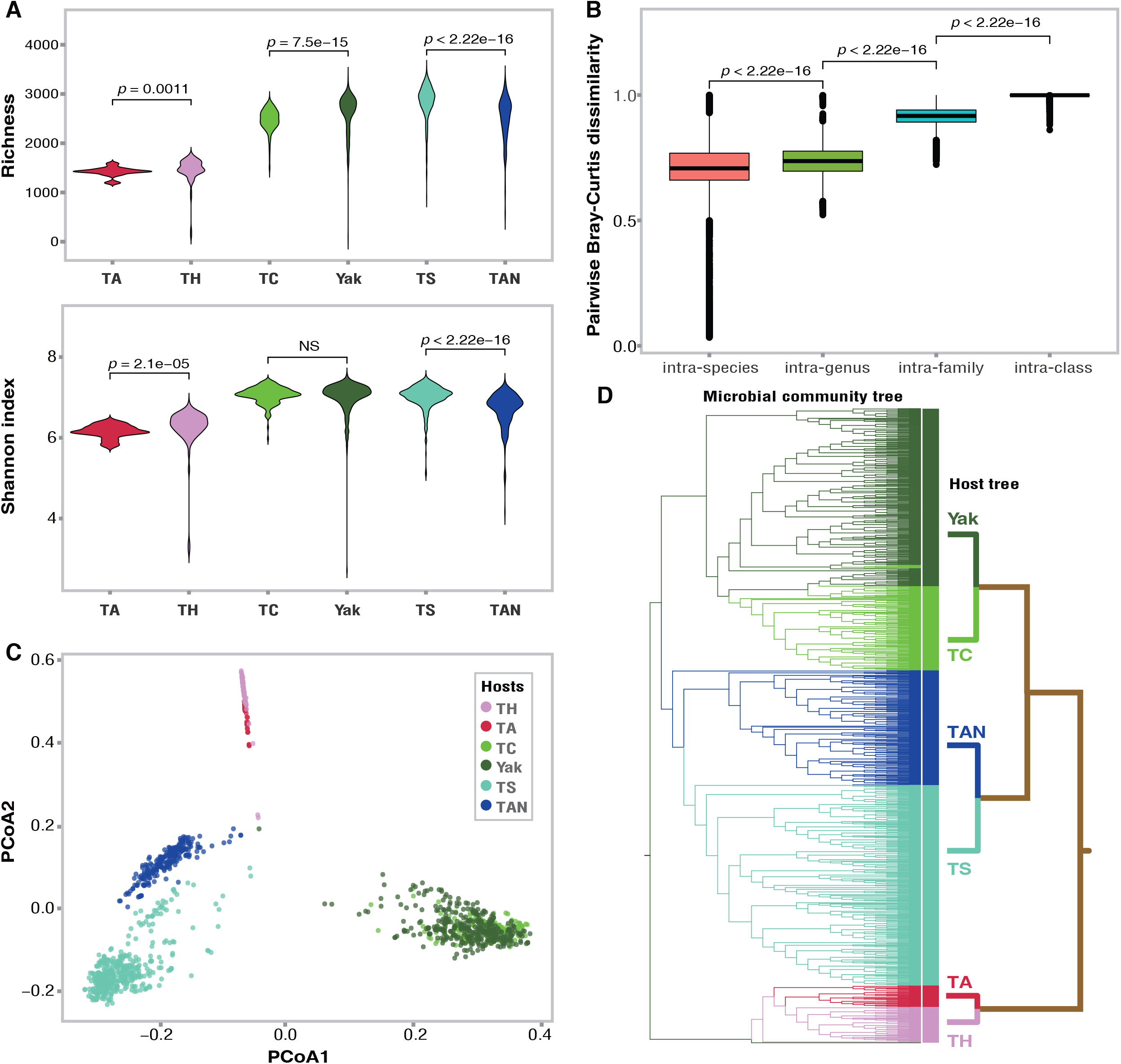
Gut microbial diversity features of the six animals host species. (**A**) Alpha diversity differences are shown by the microbial richness and Shannon index of the gut microbiomes of the six host animals. (**B**) Statistical comparison of pairwise Bray-Curtis dissimilarity of paired samples within the different taxonomic ranks for the six host animals. (**C**) PCoA plotting based on Bray-Curtis dissimilarity shows a distinct separation of gut microbial community structures among the six host animals. (**D**) The microbial community tree constructed by the Bray-Curtis dissimilarity shows the phylosymbiosis pattern with the host tree.

**Figure 4.**
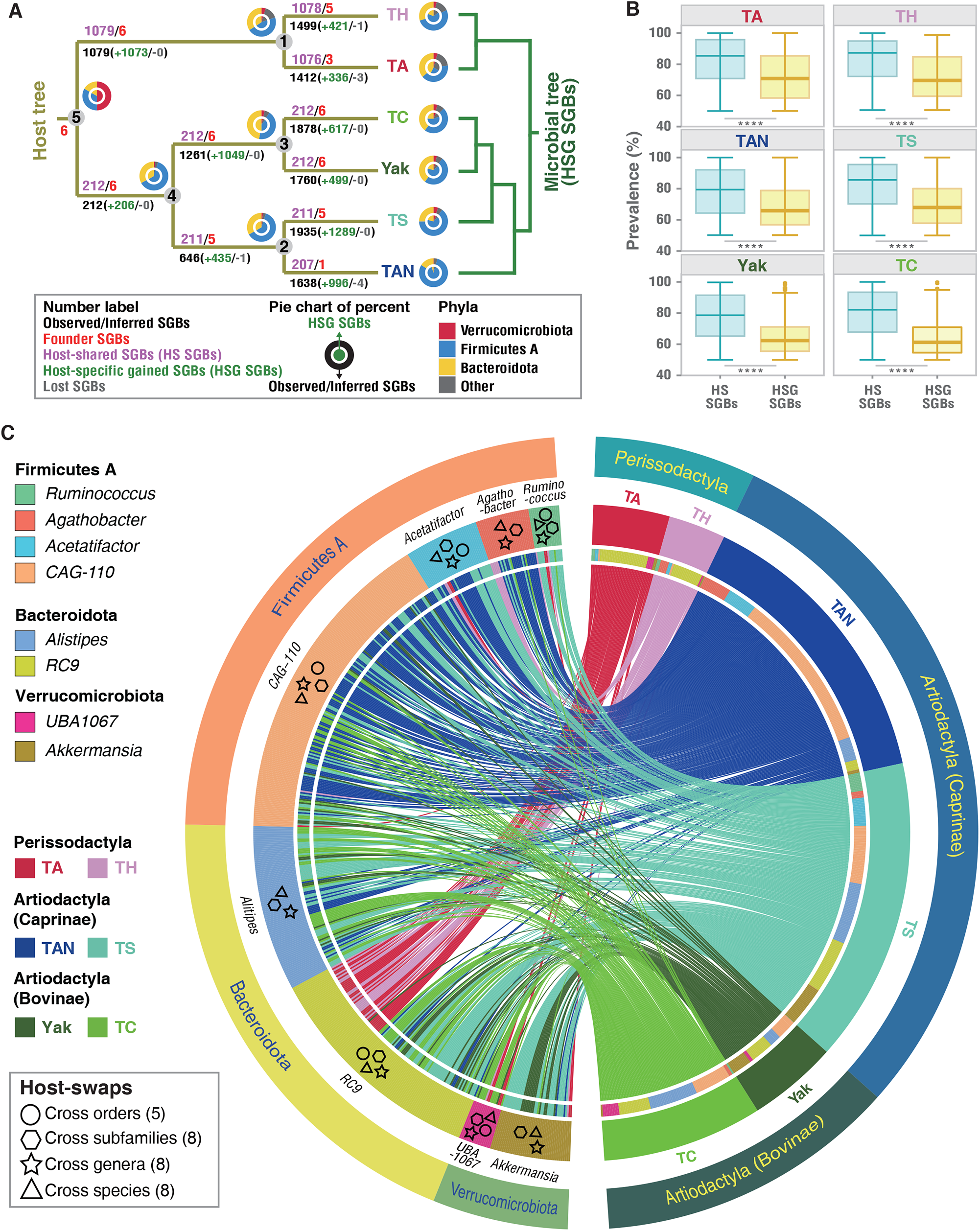
Evolutionary dynamics of gut core microbiomes. (**A**) Parsimony-inferred shifts of core gut microbiomes based on the phylogenetic relationships of host animals. The numbers on the branch stand for the SGB number of the node. The pie figure shows the microbial species composition of the node at the phylum level. Inner is for the gained SGBs and outer is for the present SGBs. The right microbial community tree was built based on specifically-gained microbial species by the six hosts, which indicates the evolutionary dynamics of gut microbial communities. HSG: Host-specific gained; HS: Host shared. (**B**) The difference of occurrence frequency by samples between host-shared and host-specific gained SGBs. (**C**) Circos representation of microbial species swapped among the six hosts. The left shows the SGBs from the eight core genera, and their phyla classifications are marked in the outer ring. The length of the ring indicates the number of the SGBs. All SGBs are sorted according to the leaf order of the SGB trees described in Supplementary Fig.7. The right is the host information. The lines in the center connect the SGBs and hosts, indicating microbial swaps among hosts shown in Supplementary Fig.7. The counts of genera with host swaps are shown in parentheses.

Our predicted six AFBs belong to the bacterial phyla Firmicutes A, Bacteroidota, and Verrucomicrobiota at the last common ancestor node (N5) (∼78 Mya) of the six host species. The lineage-specific gained SGBs were largely allocated to the three phyla indicated above, implying that these three phyla are representative phyla of the six animal hosts’ gut microbiomes. Using the AFBs as the core skaffold, we found that a stable core gut microbiome of each host was finally formed by a recuring lineage-specific gain of the microbial species, which was further supported by the phylogenetic trees of SGBs from the three representative bacterial phyla (Supplementary Fig. 6). The HS SGBs definitely played a supportive role in the common ancestor node (N1, N2, N3, N4), and the HSG SGBs were acquired and retained in the same branch. For instance, *CAG-110* from the Firmicutes A, *RC9* from the Bacteroidota and *Akkermansia* from the Verrucomicrobiota. Notably, the prevalence of HS SGBs was significantly higher than that of HSG SGBs (Fig. 4B), which may be related to the role of HS SGBs, indicating different co-evolution patterns between the two types of SGBs and their hosts. The HS SGBs might be selected by their hosts sharing a similar conserved impact from environmental sources *(i.e*., food, water, and habitat) or the close relatives of the hosts. The co-evolution or co-speciation between the HS SGBs and their hosts is impossible because the divergence times of the six animal species ranged from 2 to 78 Mya while the HS SGBs showed no host species-level divergence.

Unlike the HS SGBs, we hypothesized that the HSG SGBs might show co-evolution or co-speciation with their current six host species. To test this, we reconstructed the phylogenetic trees of the HSG SGBs representing eight genera belonging to the three representative phyla and present in at least four animal hosts (Supplementary Table 8) (Fig. 4C and Supplementary Fig. 7). We found that the phylogenetic relationships among the SGBs of each bacterial genus were inconsistent with their host phylogeny, and events of swaps occurred frequently between the hosts of different species, genera, subfamilies, and even at the order level. Transfer events were quite obvious in *Ruminococcus* and *Acetatifactor* from Firmicutes A, and UBA1067 from Verrucomicrobiota (Supplementary Fig. 7B, D and H). Thus, topological relationships inferred from the SGB trees of eight bacterial genera did not support co-speciation patterns between mammalian species and their individual gut symbionts. Conversely, the gut microbial species can span host restrictions even beyond sister order levels. We built the phylogenetic trees of strains from two bacterial species (See Supplementary Methods) showing that strain selection was not restricted between the hosts of different subfamilies (Bovinae and Caprinae) (Supplementary Fig. 8 and Supplementary Table 9). However, this conclusion needs to be confirmed by adding more bacterial species in future studies.

### Functional dynamics of the core gut microbiomes across six non-human mammals

We performed a comprehensive investigation of the metabolic capacities of the 6 AFBs based on annotation using CAZy and KEGG databases (Methods) (Supplementary Table 10). We found that the AFBs had the ability to utilize 12 common types of carbohydrates, including cellulose, hemicelluloses. The AFBs also have the potential for synthesizing acetate, propanoate, butanoate, lactate, 19 essential amino acids (AA) (except histidine), and 7 vitamins, A, B1, B2, B3, B5, B6 and B9 (Fig. 5A and Supplementary Fig. 9). These results suggest that the six AFBs core founder may play a role for host survival.

**Figure 5.**
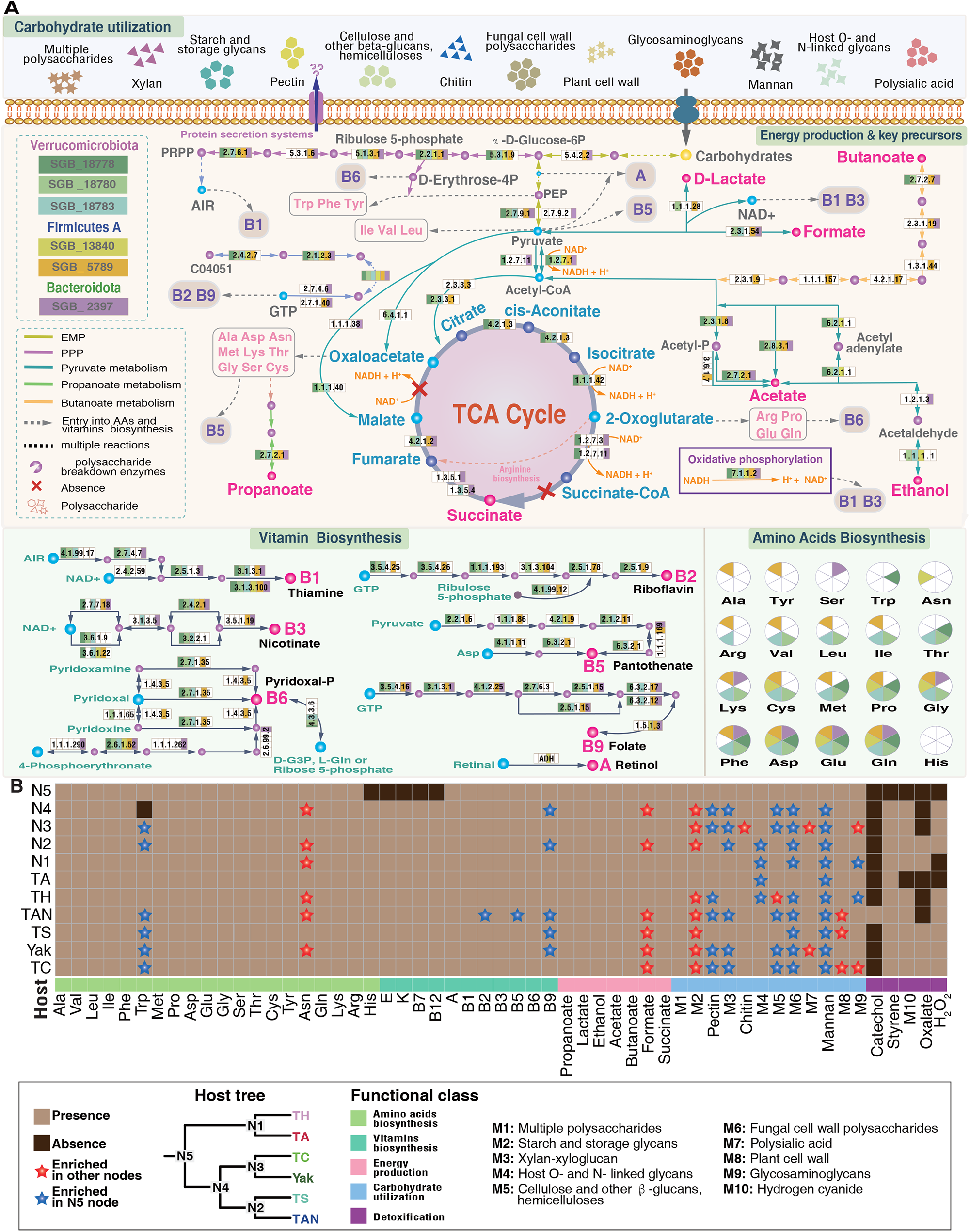
Functional profiling of six founder bacteria. (**A**) The major metabolic capacity of the six AFBs in relation to carbohydrate utilization, energy production & key precursors, amino acid biosynthesis, and vitamin biosynthesis. If the enzymes from the six AFBs can form at least one complete reaction chain, the product can theoretically be synthesized by the AFBs. (**B**) Heatmap for presence/absence and enrichment of gained SGBs of nodes compared with AFBs. Different colors indicate functional categories such as amino acid biosynthesis, vitamin biosynthesis, energy production, carbohydrate utilization, and detoxification, respectively (Supplementary Fig.9A, 9B, and 10).

To examine the evolutionary functional dynamics along host phylogeny, the functions of the HS and HSG SGBs in relation to these five core functional classes including carbohydrate utilization, energy production, amino acid biosynthesis, vitamin biosynthesis, and detoxification were surveyed. We found that the functions donated by most of the HS and HSG SGBs to a large extent overlapped with those of the AFBs (Fig. 5B and Supplementary Fig. 10). A few new functions were present at other host nodes including the biosynthesis of histidine (N4), four vitamins (E, K, B7 and B12), and the potential degradation of styrene possibly derived from plant. Only in one case, HSG SGBs at the N4 node (the common ancestor of Bovinae and Caprinae) lost the ability to synthesize tryptophan exhibiting a decreasing trend along this branch. Furthermore, our enrichment analyses revealed that there was no significant difference in functional enrichment between AFBs and HSG SGBs in most terms of biosynthetic ability. Overall, our results suggested that the HSG SGBs maybe contributed to maintaining the stability of the functions of the AFBs on these core functional classes.

Our results (Fig. 6) also revealed that the HSG SGBs exhibited significant functional divergences in relation to carbohydrate utilization and main metabolic pathways between N1 vs. N4 (∼78 Mya), N2 vs. N3 (∼20 Mya), or the comparisons among related host species (TA vs. TH, TAN vs. TS and Yak vs. TC). With four exceptions, tyrosine metabolism, aminobenzoate degradation, bispenol degradation, and furfural degradation, significant functional divergences were observed among carbohydrate utilization and metabolic pathways in four KEGG functional categories that may be related to the evolutionary adaptation of each host (Fig. 6; Supplementary Fig. 11; Supplementary Table 12).

**Figure 6.**
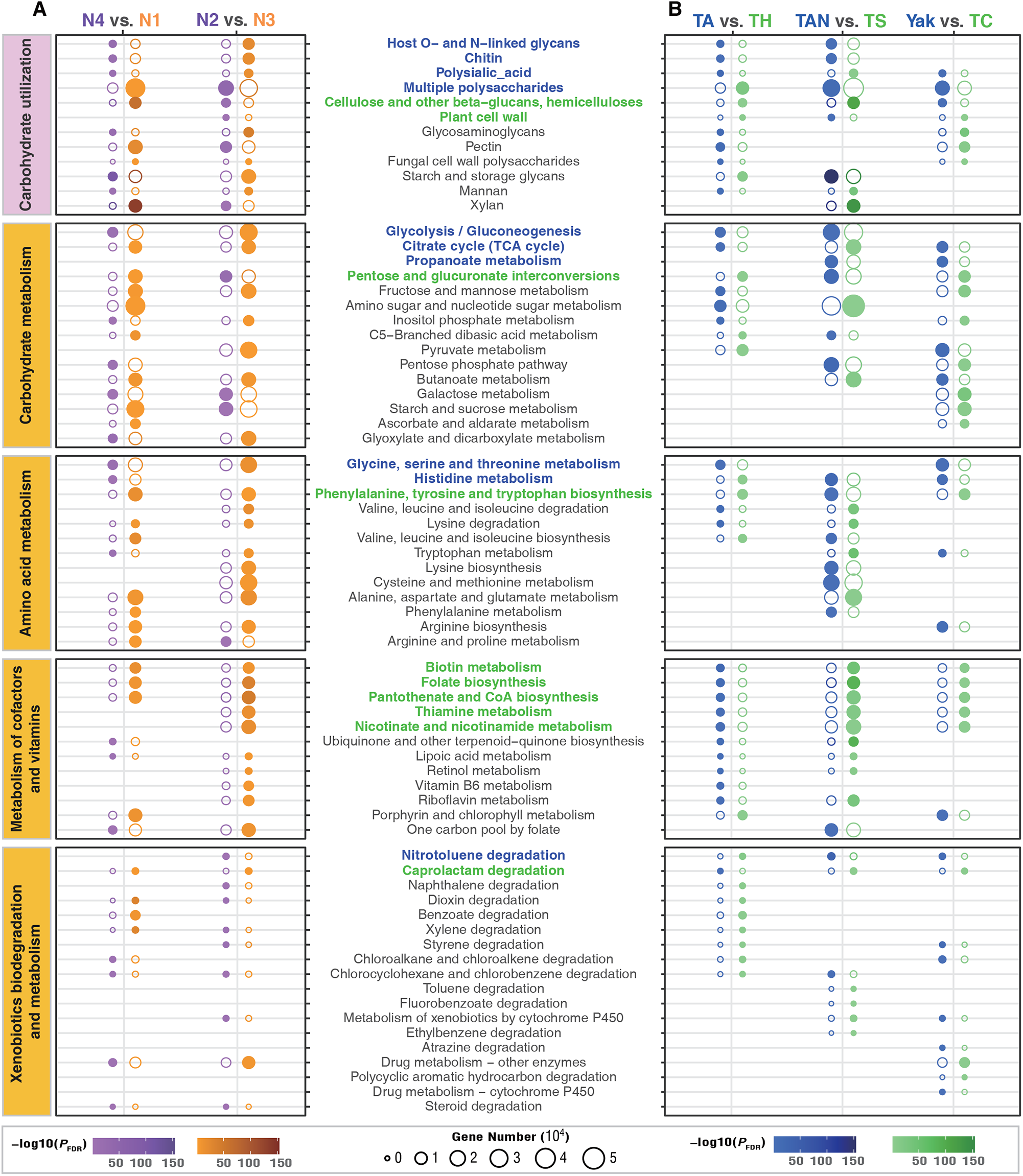
The functional divergence of gained SGBs among key evolutionary nodes. (**A**) Functional enrichment analysis among internal evolution nodes, including those between Perissodactyla (N1) and Artiodactyla (N4), Bovinae (N3) and Caprinae (N2). The size of the circle represents the number of genes in the substrate utilization capacities or the metabolic pathways. The solid circle represents the enriched node. The color scale from light to dark corresponds to the negative logarithm of false discovery rate (FDR)-adjusted P values (<0.05) from low to high. (**B**) Comparison of the enrichment of gut microbial functions between plateau indigenous species and late-migratory species. The size of the circle represents the number of genes in the substrate utilization capacities or the metabolic. The solid circle represents the enriched animal host. The color scale from light to dark corresponds to the negative logarithm of FDR-adjusted P values (< 0.05) from low to high. The strips on the left represent different functional pathway classifications. The pink strip indicates the substrate utilization capacity based on the CAZy database annotation. The yellow strips indicate pathway classifications based on KEGG database. The y-axis labels in the middle of the figure indicate different substrate utilization capacities or pathways. Blue colors indicate enrichment in at least 2 indigenous species and green colors indicate enrichment in at least 2 late-migrated species.

We investigated the differences at the functional level between long-term-adaptation and short-term-adaptation of the host species and pathway enrichment (Fig. 6B). These analyses revealed that multiple metabolic pathways were consistently enriched in the two indigenous plateau artiodactyla mammals, such as the ‘multiple polysaccharides’ in carbohydrate utilization, ‘propanoate metabolism’, ‘histidine metabolism’ and ‘nitrotoluence degradation’ pathways. In addition, convergent enrichments of pathways were also observed in two short-term adapted artiodactyla mammals, such as five pathways involved in metabolism of cofactors and vitamins, and the caprolactam degradation pathway. We also found host-specific enrichment of metabolic pathways provided by the HSG SGBs. For example, the pathways involved in arginine biosynthesis were only enriched in yak. The pathway ‘vitamin B6 metabolism only appeared in Tibetan Ass. Similarly, lysine biosynthesis, cysteine and methionine metabolism, and phenylalanine metabolism were unique for Tibetan Antelope.

## Discussion

We present a large *de novo* microbial genome assembly from metagenomic data providing 19,251 gut microbial species-level reference genomes derived from six non-human mammals of the Qinghai-Tibet Plateau. Of these species > 99% are unknown, thus expanding the known phylogenetic diversity of bacteria and archaea by 62.40% and 10.29%, respectively, compared with the GTDB database *(31)*. Over the past two decades, large-scale studies of the human gut microbiome have provided a comprehensive catalog of human gut microbial species reference genomes comprising 204,938 non-redundant genomes from 4,644 gut prokaryotes of which more than 70% lack cultured representatives *(7)*. The Earth’s microbiomes project (EMP) discovered 52,515 MAGs representing 12,556 novel candidate species spanning 135 phyla expanding the known phylogenetic diversity of bacteria and archaea *(33)*. Even though 1,209 SGBs (75% unknown) was recently unveiled from 406 fecal samples from 184 animal species *(8)*, we discovered unexpectedly a large number of novel bacterial and archaeal genomes ornamenting the first blueprint of gut microbiomes of native mammals at the *third pole(9)*, implying that previous global research greatly has underestimated the (gut) microbial diversity in non-human mammals, and a considerable number of unknown microbial species still need to be uncovered by global efforts to elucidate their biological roles in various environmental niches.

Despite our findings to a certain extent supported the notion that gut microbial community relationships parallel the phylogeny of their hosts, coined phylosymbiosis*(19, 38)*, our whole-genome-level phylogenetic analyses revealed that lineage-specifically gained microbial species were frequently transferred across host species, genus, subfamily, and even order levels. These findings, like previous studies, did not support the existence of co-phylogeny or co-speciation events between mammal hosts and their gut individual symbionts *(13-16)*. This discrepancy is probably mainly due to the low resolution and likely lateral transfers of partial 16S rRNA genes *(20)* or single-copy marker genes*(21)* compared with the whole-genome phylogenetic analysis for accurately obtaining phylogenetic relationships among gut microbial species. Similarly, many previous theoretical and experimental studies demonstrated that short-term dynamics can foster parasite specialization, but that these events can occur following host shifts and do not necessarily involve co-speciation, as well as coevolutionary dynamics of hosts and parasites do not favor long-term cospeciation *(38)*.

Our study enables a glimpse into diverse functional traits of mammalian gut microbiomes. We obtained a huge non-redundant gene catalog containing 26,093,065 genes, among which 82.53% and 66.43% were annotated by the Nr database and KEGG database, respectively. Additionally, from 3888 (69.34%) of 5607 SGBs of the three core bacterial phyla (Firmicute A, Bacteroidota, and Verrucomicrobiota), we identified a total of 9,221 biosynthetic gene clusters (BGCs) (consisting of 130,098 intact CDSs or genes), 9,218 of which could be assigned into 60 known BGC types (Supplementary Fig. 12 and Supplementary Table 13), mostly involved in the biosynthesis of many secondary metabolites like carotenoid affecting host physiology or health. These findings suggest our assembled SGBs also represent a large natural gene pool which requires further exploration.

Lineage-specific gained microbial species might endow host adaptation to hypoxia environment. Tibetan ass, Tibetan antelope, and yak that have long-term adaptation to hypoxia *(39, 40)* and some of the enriched metabolic pathway may assist. Thus, vitamins B6, B12, folate, and choline are reported to elicit combined neuroprotective effects on the brain against hypoxia *(41)*. Riboflavin requirement is increased under acute hypoxic conditions and its supplementation can improve energy metabolism *(42)*. Cysteine supplementation allows the body to respond to and adapt to hypoxic situations more quickly *(43)*. Overall, these findings provided indirect support for the hypothesis that the distinct gut microbiomes found in high-altitude mammals may be linked to high-altitude hypoxia adaption. More research will be needed to better understand the biological significance of these discoveries.

## Acknowledgements

Thanks for supports for sample collections by Zewang Jiangcun, Dunzhu Silang, Zerowang Taba, Ding Deng, and staff members from the Qiangtang National Nature Reserve, Tibet, China, the Science and Technology Department of Tibet. Lhasa, Tibet, China, as well as the Hoh Xil Nature Reserve, Qinghai, China, the Sanjiangyuan National Park, Qinghai, China, and the Science and Technology Department of Qinghai, Qinghai, China. Thanks to Zhang Cheng-Hao and Zewang Jiangcun for the animal pictures. Thanks for suggestions for the assembly of MAGs from Zhuye Jie and Tao Zhang. We gratefully acknowledge colleagues at BGI-Shenzhen for discussions. This study was supported by the Second Tibetan Plateau Scientific Expedition and Research (STEP) program (no. 2019QZKK0503), the Chinese National Natural Science Foundation (no. U2002206 and 31970571), and the Major Science and Technology Project in Yunnan Province of China (no. 202001BB050001).

## Author contributions

Z.Z. conceived, designed, and supervised the study. Z.Z., W.Q., T.C., Z.D., Z.F., L.X.T, Z.T., and S.J. collected samples. L.X., L.T., T.C., Z.D., B.Y., G.H., X.S. and Z.H. processed and analyzed the raw sequencing data. Z.Z. and L.X. interpreted the data. L.X., Z.Z., T.C., Z.D, B.Y., G.H., X.S., M.D and Z.H. generated the figures. L.X., Z.Z., T.C. and Z.D. wrote the manuscript. Z.Z., L.X., S.K.S. and K.K. contributed to critical revision. All authors contributed to the final manuscript.

## Data Availability

All the clean data, recovered genomes, SGBs and SGBs profiles, the GenBank format annotations and CDS sequences of 9221 BGCs of this study have been deposited into CNGB Sequence Archive (CNSA) of China National GeneBank DataBase (CNGBdb) with accession number CNP0001390 (https://db.cngb.org/search/project/CNP0001390/). The raw sequencing data of the 1,412 samples are also available in https://db.cngb.org/qtp/.

## Code Availability

A repository containing instructions to reproduce the analyses is available at https://github.com/QTP-team. Freely available software and package was excluded from this repository.

## REFERENCES

1. Z. Zhang et al., Convergent Evolution of Rumen Microbiomes in High-Altitude Mammals. Current Biology 26, 1873–1879 (2016).

2. P. Rosshart Stephan et al., Laboratory mice born to wild mice have natural microbiota and model human immune responses. Science 365, eaaw4361 (2019).

3. C. Campbell et al., Bacterial metabolism of bile acids promotes generation of peripheral regulatory T cells. Nature 581, 475–479 (2020).

4. S. K. Gill, M. Rossi, B. Bajka, K. Whelan, Dietary fibre in gastrointestinal health and disease. Nature Reviews Gastroenterology & Hepatology 18, 101–116 (2021).

5. P. Kundu, E. Blacher, E. Elinav, S. Pettersson, Our Gut Microbiome: The Evolving Inner Self. Cell 171, 1481–1493 (2017).

6. E. Pasolli et al., Extensive Unexplored Human Microbiome Diversity Revealed by Over 150,000 Genomes from Metagenomes Spanning Age, Geography, and Lifestyle. Cell 176, 649–662.e620 (2019).

7. A. Almeida et al., A unified catalog of 204,938 reference genomes from the human gut microbiome. Nature Biotechnology 39, 105–114 (2021).

8. D. Levin et al., Diversity and functional landscapes in the microbiota of animals in the wild. Science 372, eabb5352 (2021).

9. J. Qiu, China: The third pole. Nature 454, 393–396 (2008).

10. Z. Zhou, T. Deng, The Tibetan Plateau is a natural laboratory for studying organic evolution and environmental change. Science China Earth Sciences 63, 169–171 (2020).

11. T. Deng, F. Wu, Z. Zhou, T. Su, Tibetan Plateau: An evolutionary junction for the history of modern biodiversity. Science China Earth Sciences 63, 172–187 (2020).

12. T. Deng et al., Out of Tibet: Pliocene Woolly Rhino Suggests High-Plateau Origin of Ice Age Megaherbivores. Science 333, 1285–1288 (2011).

13. L. Cortes-Ortiz, K. R. Amato, Host genetics influence the gut microbiome. Science 373, 159–160 (2021).

14. A. H. Moeller, T. A. Suzuki, M. Phifer-Rixey, M. W. Nachman, Transmission modes of the mammalian gut microbiota. Science 362, 453–457 (2018).

15. L. Grieneisen et al., Gut microbiome heritability is nearly universal but environmentally contingent. Science 373, 181–186 (2021).

16. P. Ferretti et al., Mother-to-Infant Microbial Transmission from Different Body Sites Shapes the Developing Infant Gut Microbiome. Cell Host & Microbe 24, 133–145.e135 (2018).

17. R. M. Brucker, S. R. Bordenstein, The roles of host evolutionary relationships (genus: Nasonia) and development in structuring microbial communities. Evolution 66, 349–362 (2012).

18. M. Groussin, F. Mazel, E. J. Alm, Co-evolution and Co-speciation of Host-Gut Bacteria Systems. Cell Host & Microbe 28, 12–22 (2020).

19. F. Mazel et al., Is Host Filtering the Main Driver of Phylosymbiosis across the Tree of Life? mSystems 3, e00097–00018 (2018).

20. M. Groussin et al., Unraveling the processes shaping mammalian gut microbiomes over evolutionary time. Nature Communications 8, 14319 (2017).

21. H. Moeller Andrew et al., Cospeciation of gut microbiota with hominids. Science 353, 380–382 (2016).

22. A. Gaulke Christopher et al., Ecophylogenetics Clarifies the Evolutionary Association between Mammals and Their Gut Microbiota. mBio 9, e01348–01318 (2018).

23. N. D. Youngblut et al., Host diet and evolutionary history explain different aspects of gut microbiome diversity among vertebrate clades. Nature Communications 10, 2200 (2019).

24. J. R. Brown, C. J. Douady, M. J. Italia, W. E. Marshall, M. J. Stanhope, Universal trees based on large combined protein sequence data sets. Nat Genet 28, 281–285 (2001).

25. F. Delsuc, H. Brinkmann, H. Philippe, Phylogenomics and the reconstruction of the tree of life. Nature Reviews Genetics 6, 361–375 (2005).

26. H. Jonsson et al., Speciation with gene flow in equids despite extensive chromosomal plasticity. Proc Natl Acad Sci U S A 111, 18655–18660 (2014).

27. Y. Jiang et al., The sheep genome illuminates biology of the rumen and lipid metabolism. Science 344, 1168–1173 (2014).

28. L. Chen et al., Large-scale ruminant genome sequencing provides insights into their evolution and distinct traits. Science 364, eaav6202 (2019).

29. A. M. Humphreys, T. G. Barraclough, The evolutionary reality of higher taxa in mammals. Proceedings. Biological sciences 281, 20132750 (2014).

30. D. H. Parks et al., Recovery of nearly 8,000 metagenome-assembled genomes substantially expands the tree of life. Nature Microbiology 2, 1533–1542 (2017).

31. D. H. Parks et al., GTDB: an ongoing census of bacterial and archaeal diversity through a phylogenetically consistent, rank normalized and complete genome-based taxonomy. Nucleic acids research 50, D785–D794 (2022).

32. R. D. Stewart et al., Compendium of 4,941 rumen metagenome-assembled genomes for rumen microbiome biology and enzyme discovery. Nature Biotechnology 37, 953–961 (2019).

33. S. Nayfach et al., A genomic catalog of Earth’s microbiomes. Nature Biotechnology 39, 499–509 (2021).

34. L. Glendinning, B. Genç, R. J. Wallace, M. Watson, Metagenomic analysis of the cow, sheep, reindeer and red deer rumen. Scientific Reports 11, 1990 (2021).

35. F. Xie et al., An integrated gene catalog and over 10,000 metagenome-assembled genomes from the gastrointestinal microbiome of ruminants. Microbiome 9, 137 (2021).

36. M. Csurös, Count: evolutionary analysis of phylogenetic profiles with parsimony and likelihood. Bioinformatics (Oxford, England) 26, 1910–1912 (2010).

37. K. Kwong Waldan et al., Dynamic microbiome evolution in social bees. Science Advances 3, e1600513 (2017).

38. D. M. de Vienne et al., Cospeciation vs host-shift speciation: methods for testing, evidence from natural associations and relation to coevolution. New Phytologist 198, 347–385 (2013).

39. Q. Qiu et al., The yak genome and adaptation to life at high altitude. Nat Genet 44, 946–949 (2012).

40. R.-L. Ge et al., Draft genome sequence of the Tibetan antelope. Nat Commun 4, 1858 (2013).

41. L. Yu, Y. Chen, W. Wang, Z. Xiao, Y. Hong, Multi-Vitamin B Supplementation ReversesHypoxia-Induced Tau Hyperphosphorylation and Improves Memory Function in Adult Mice. Journal of Alzheimer’s Disease 54, 297–306 (2016).

42. Y. P. Wang et al., Riboflavin supplementation improves energy metabolism in mice exposed to acute hypoxia. Physiol Res 63, 341–350 (2014).

43. S. C. Nunes et al., Cysteine boosters the evolutionary adaptation to CoCl2 mimicked hypoxia conditions, favouring carboplatin resistance in ovarian cancer. BMC Evolutionary Biology 18, 97 (2018).

