## Supplementary figures for "A Unified Catalog of 19,251 Non-human Reference Species Genomes Provides New Insights into the Mammalian Gut Microbiomes"

**Supplementary Fig. 1**

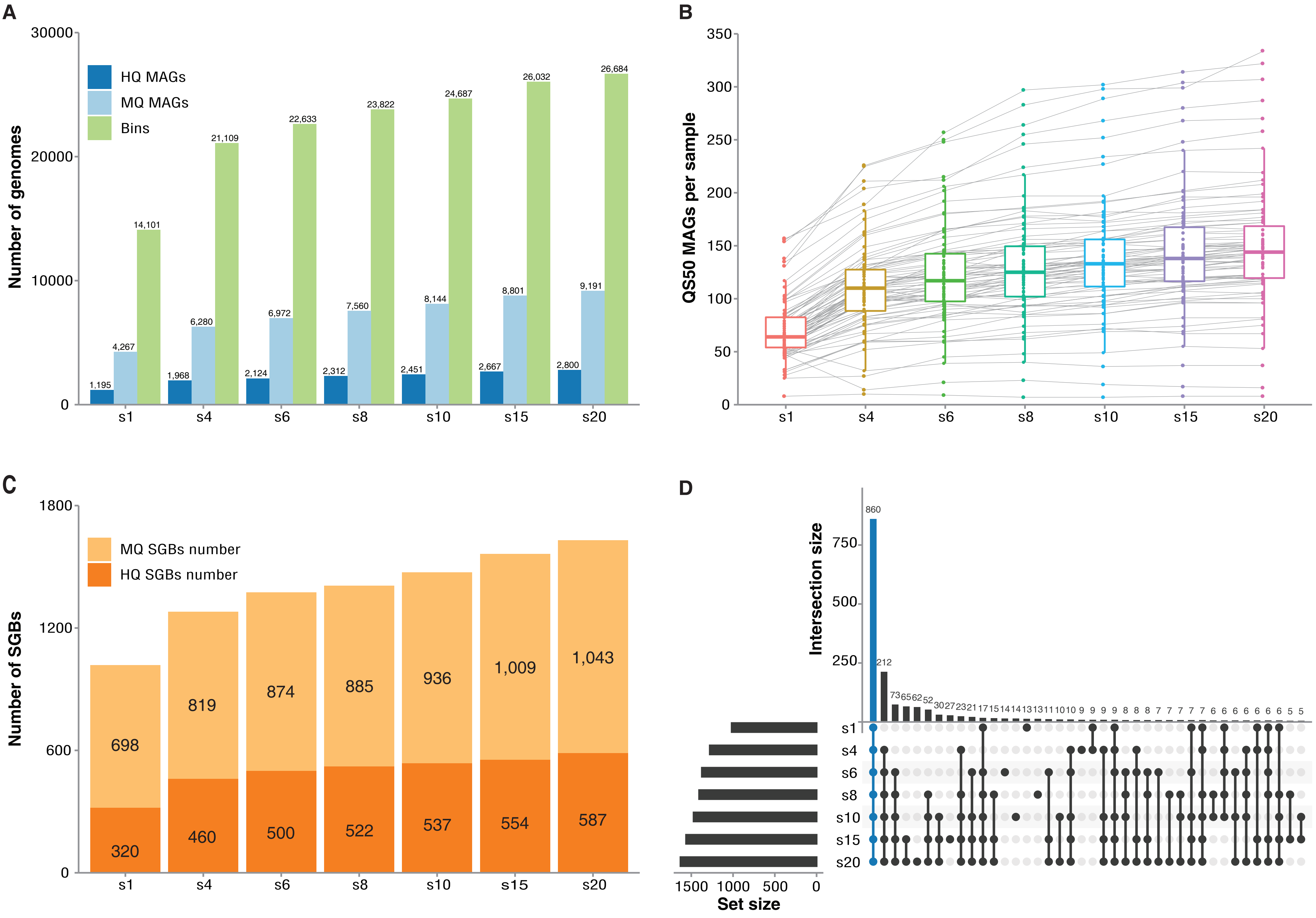

#### Supplementary Fig. 2

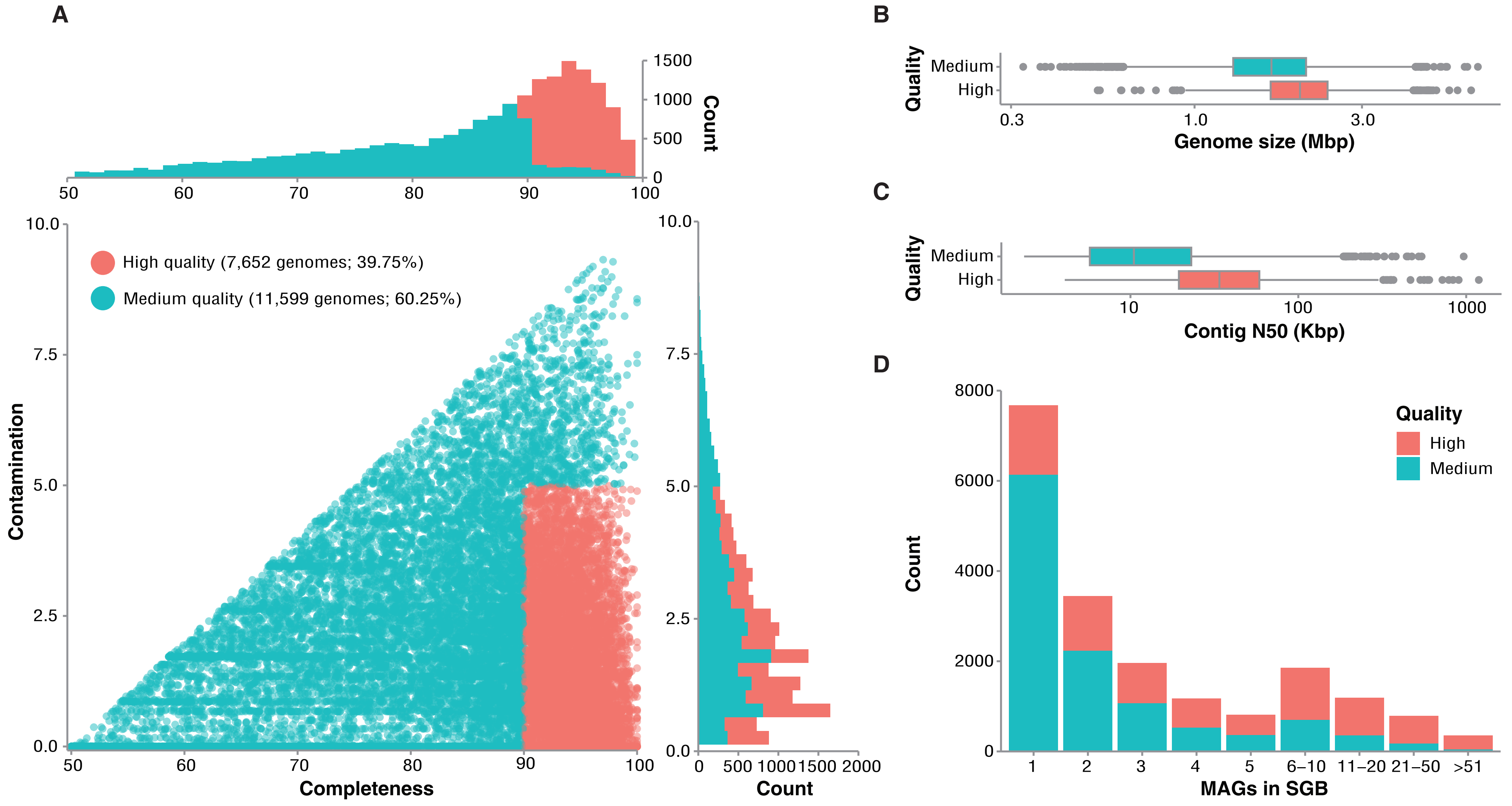

Supplementary Fig. 3

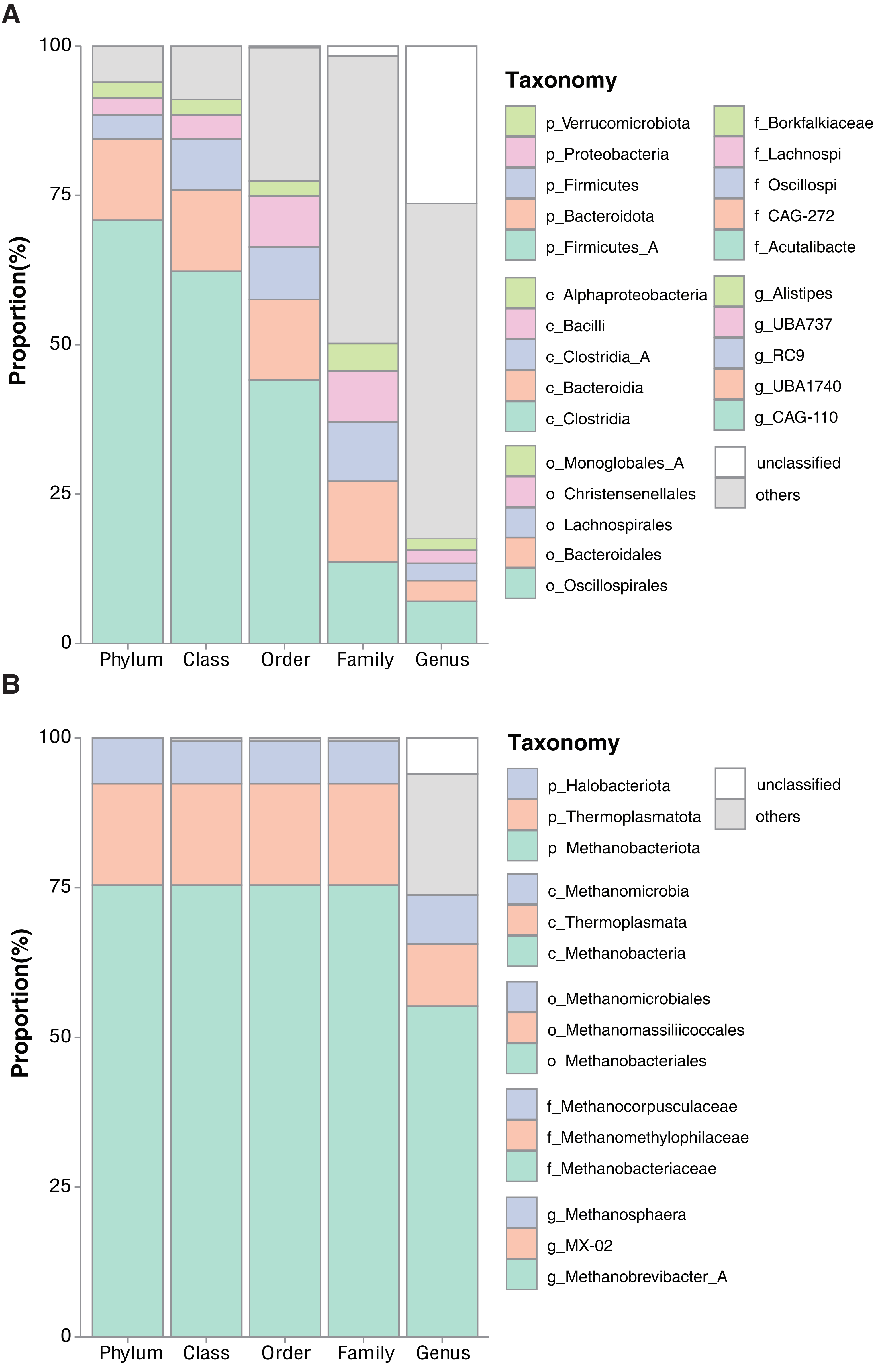

Supplementary Fig. 4

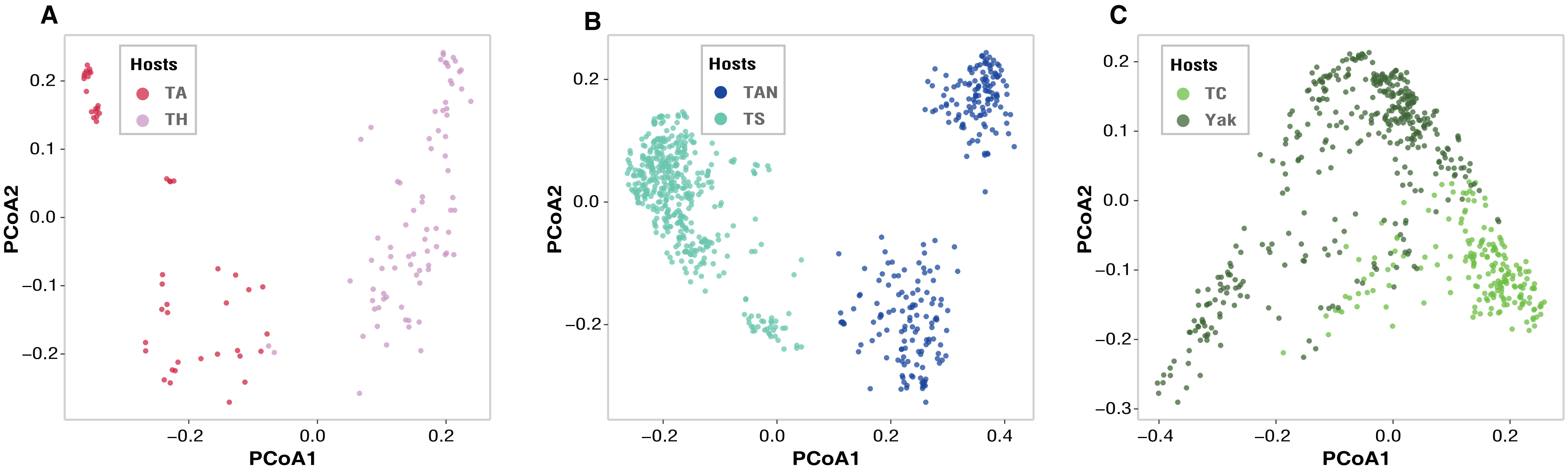

Supplementary Fig. 5

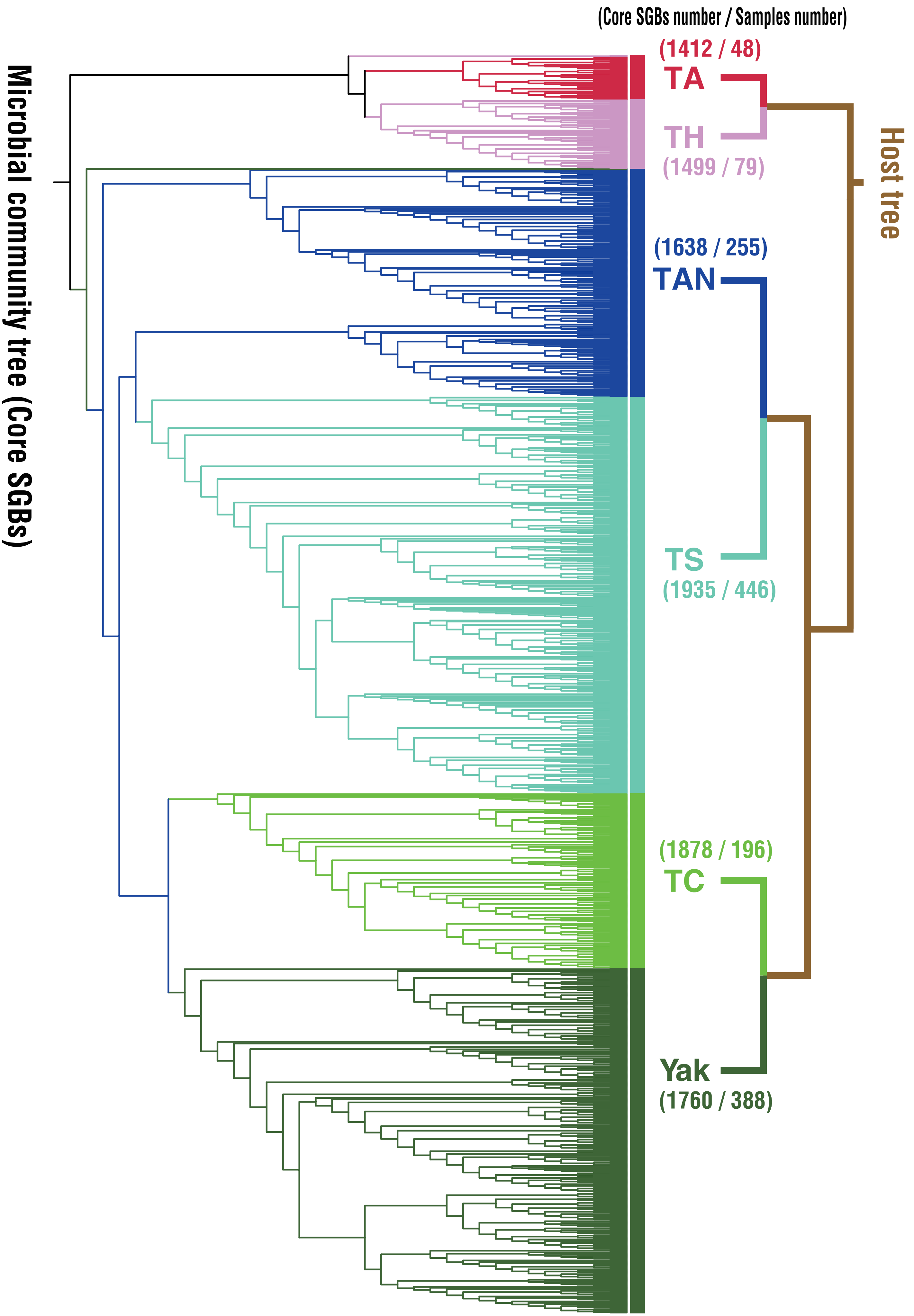

#### Supplementary Fig. 6

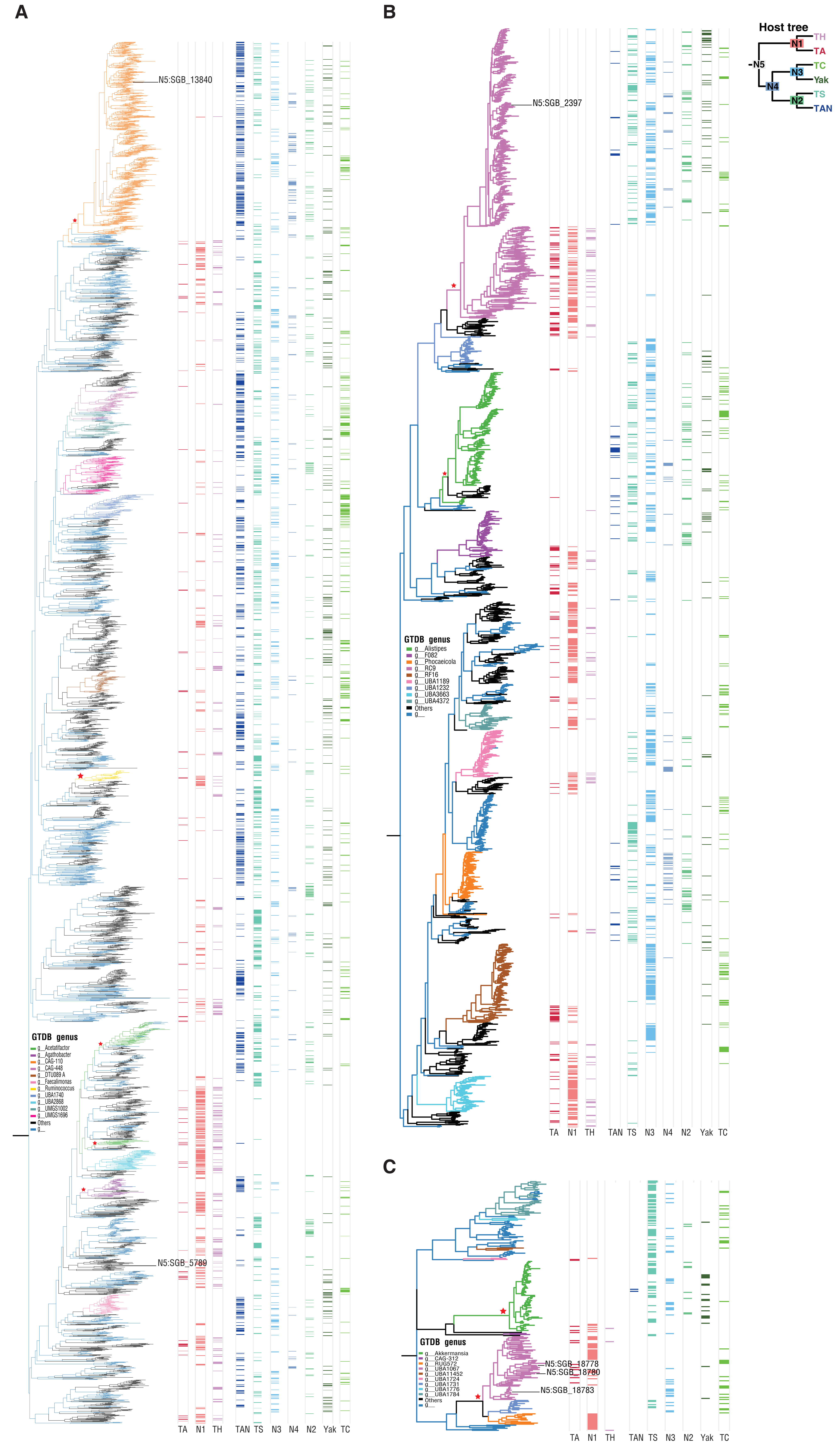

**Supplementary Fig. 7**

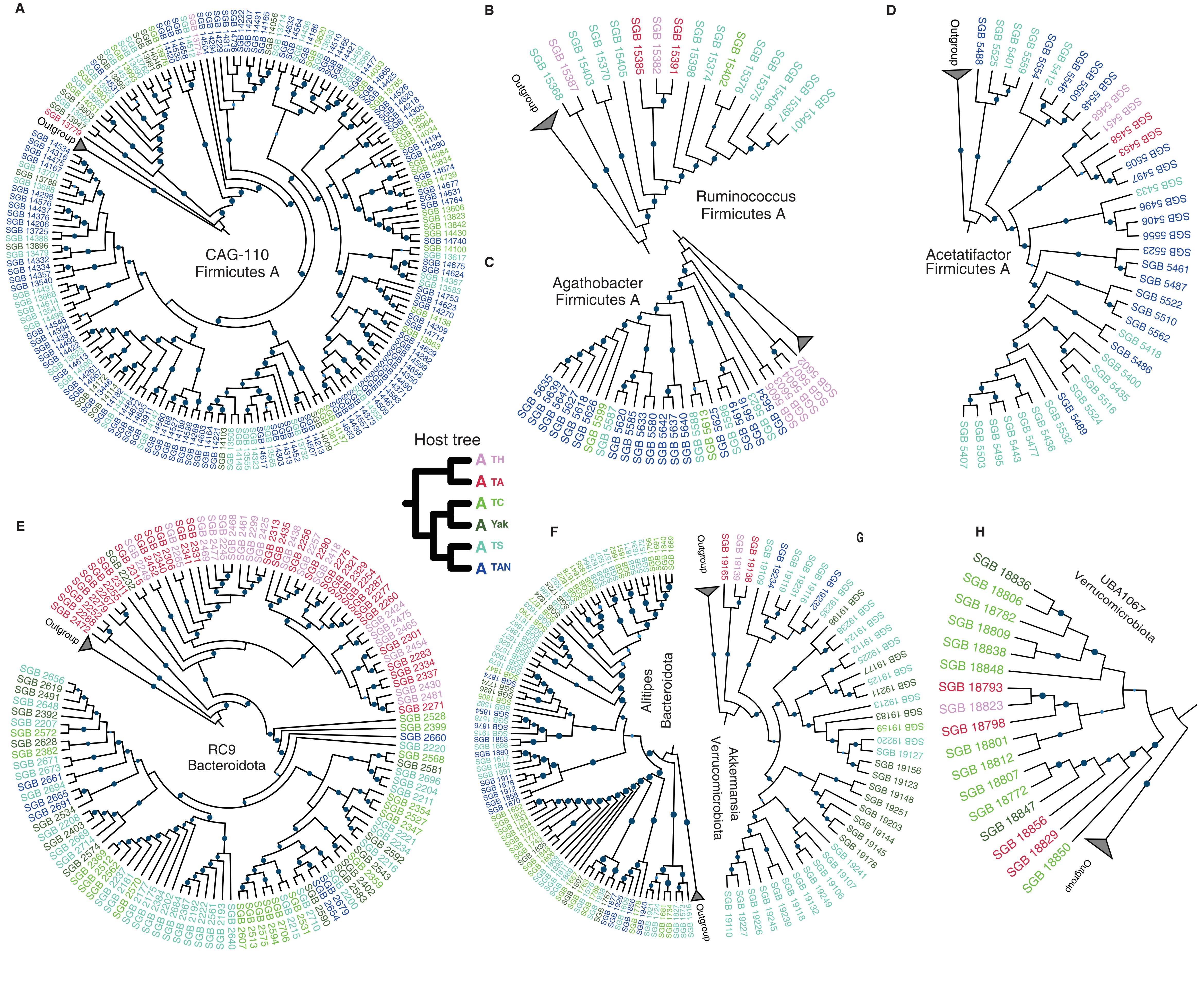

**Supplementary Fig. 8**

A

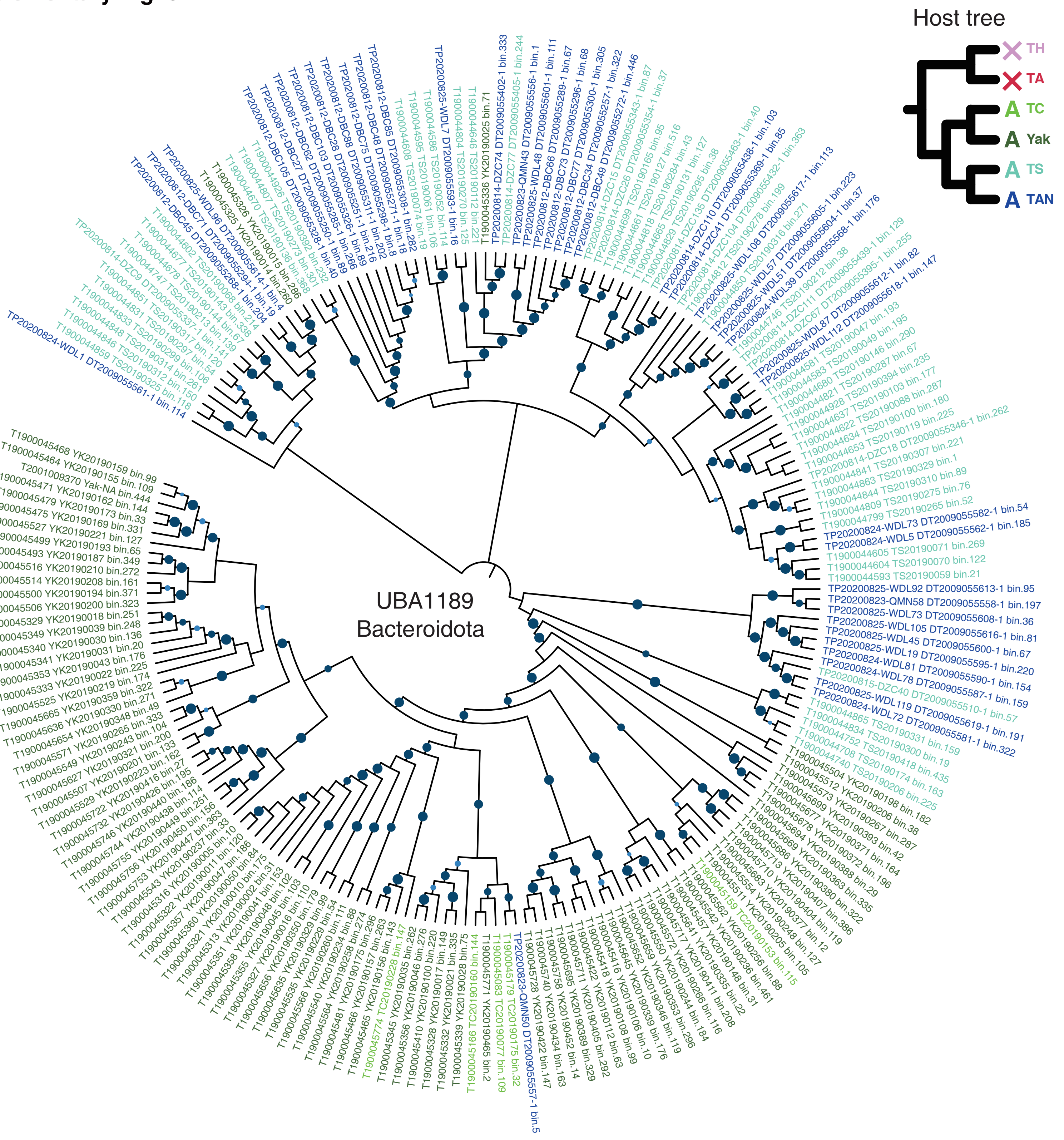


B

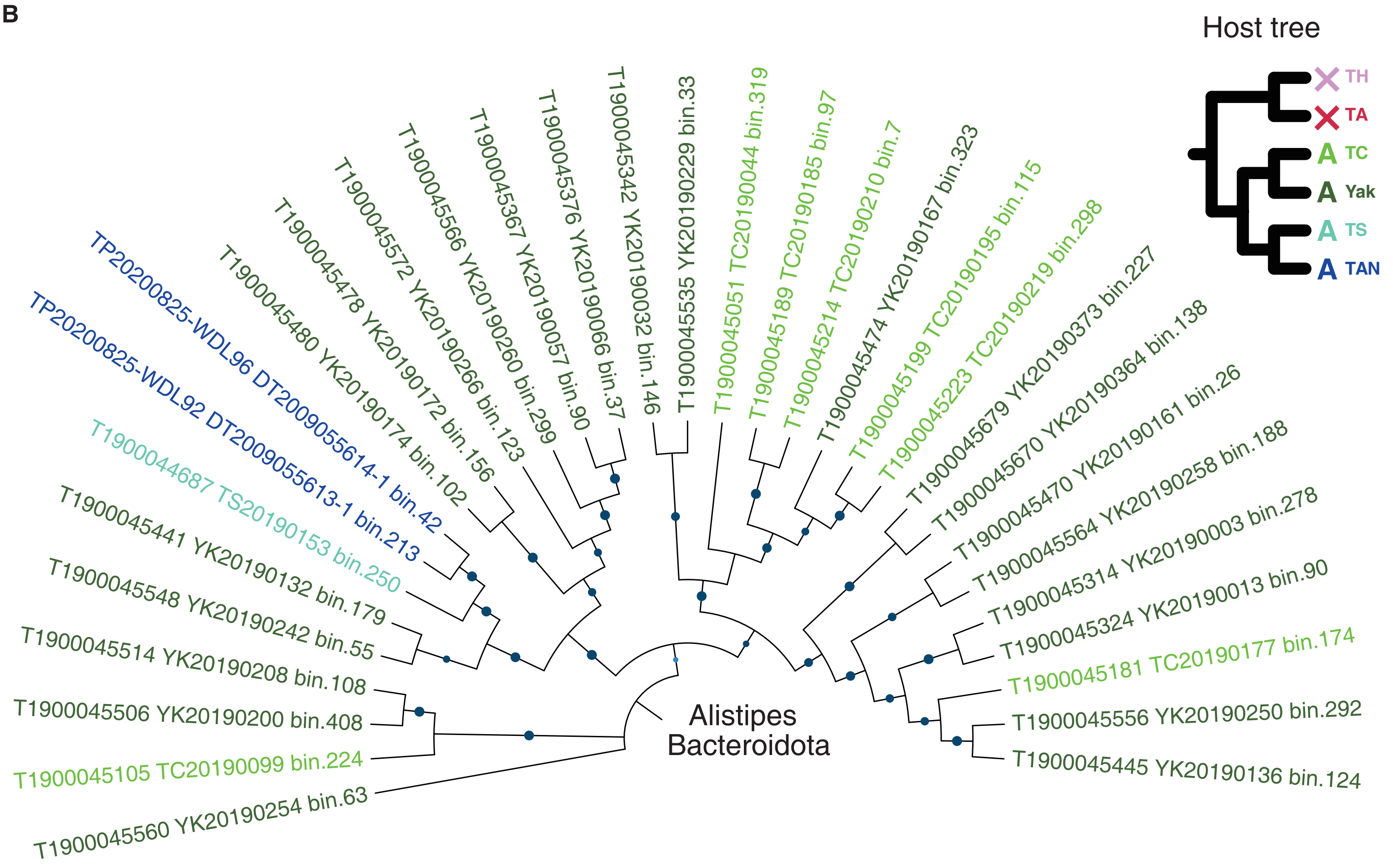

**Supplementary Fig. 9**

#### Amino Acid biosynthesis

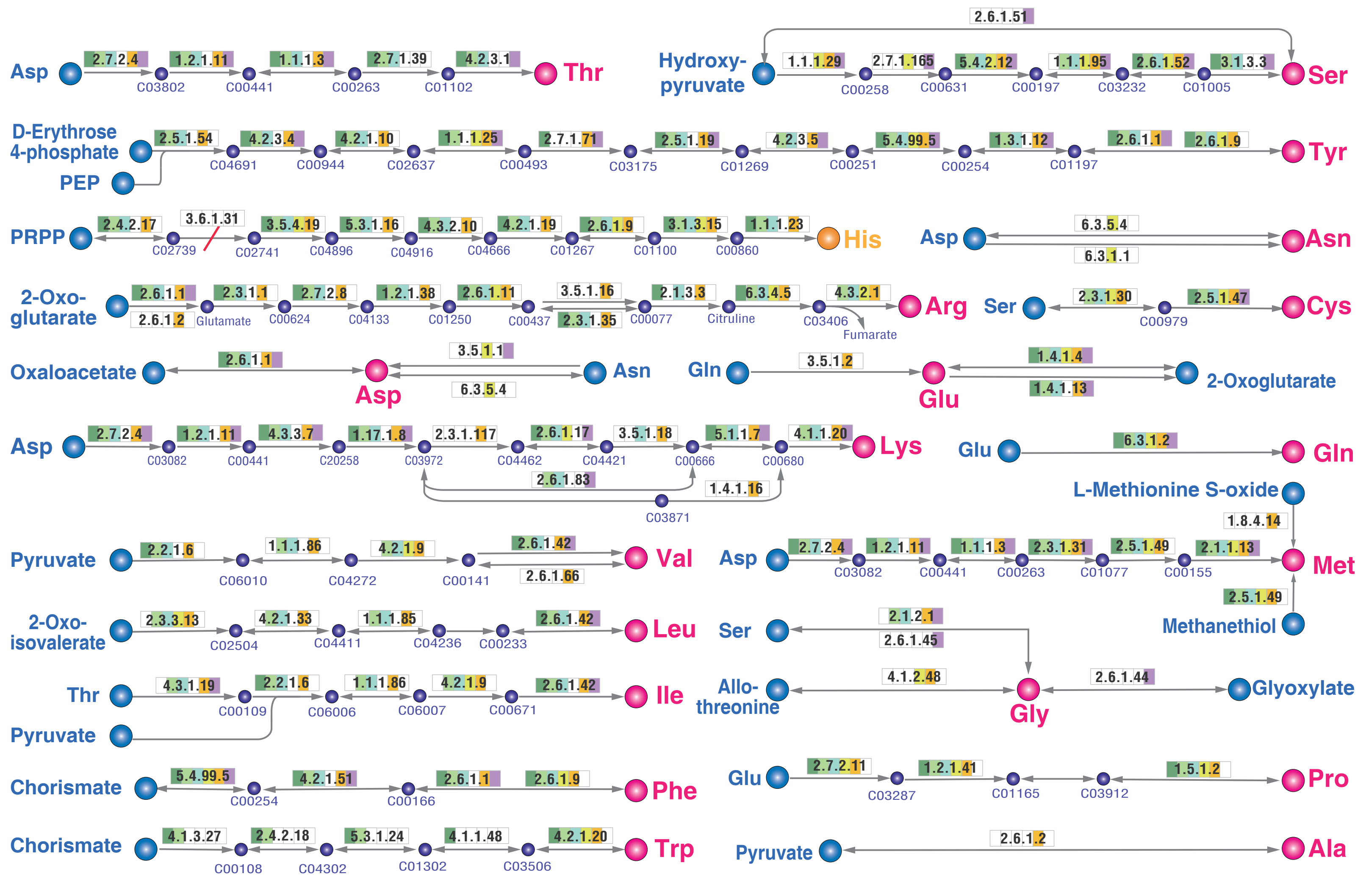


**Phylum** | Verrucomicrobiota SGB_18783 SGB_18780 SGB_18778 | Bacteroidota SGB_2397 | Firmicutes A SGB_13840 SGB_5789

**Reaction** | Initial Substrates | End products | Intermediate products | Non-synthesize | Absent step

**Enzyme ECs & Definitions:**

2.5.1.47: cysteine synthase
4.1.2.48: threonine aldolase
3.5.1.1: L-asparaginase
2.2.1.6: acetolactate synthase
2.7.2.11: glutamate 5-kinase
2.7.2.4: aspartate kinase
6.3.5.4: asparagine synthase B
5.4.99.5: chorismate mutase
6.3.1.2: glutamine synthetase
2.6.1.2: alanine transaminase
2.7.2.8: acetylglutamate kinase
3.1.3.15: histidinol-phosphatase
4.2.1.51: prephenate dehydratase
1.1.1.23: histidinol dehydrogenase
4.3.1.19: threonine ammonia-lyase
4.2.1.35: (R)-2-methylmalate dehydratase
1.5.1.2: pyrroline-5-carboxylate reductase
2.6.1.66: valine--pyruvate aminotransferase
2.4.2.18: anthranilate phosphoribosyltransferase
5.3.1.24: phosphoribosylanthranilate isomeraseS
2.6.1.9: histidinol-phosphate aminotransferase
1.4.1.13: glutamate synthase (NADPH) large chain
1.2.1.11: aspartate-semialdehyde dehydrogenase
4.3.3.7: 4-hydroxy-tetrahydrodipicolinate synthase

2.5.1.49: O-acetylhomoserine (thiol)-lyase
4.1.1.20: diaminopimelate decarboxylase
1.4.1.16: diaminopimelate dehydrogenase
2.4.2.17: ATP phosphoribosyltransferase
4.4.1.13: cysteine-S-conjugate beta-lyase
2.1.2.1: glycine hydroxymethyltransferase
4.2.1.20: tryptophan synthase alpha chain
2.6.1.11: acetylornithine aminotransferase
2.3.1.35: glutamate N-acetyltransferase
3.5.1.49: phosphoribosyl-AMP cyclohydrolase
4.3.2.10: imidazole glycerol-phosphate synthase
4.1.1.48: indole-3-glycerol-phosphate synthase
2.6.1.83: LL-diaminopimelate aminotransferase
1.4.1.4: glutamate dehydrogenase (NADP+)

4.2.1.9: dihydroxy-acid dehydratase
1.1.1.86: ketol-acid reductoisomerase
2.3.1.182: (R)-citramalate synthase
2.3.3.13: 2-isopropylmalate synthase
6.3.4.5: argininosuccinate synthase
2.3.1.30: serine O-acetyltransferase
2.3.1.1: N-acetylglutamate synthase
6.3.1.1: aspartate--ammonia ligase
1.3.1.12: prephenate dehydrogenase
2.6.1.1: aspartate aminotransferase
5.1.1.7: diaminopimelate epimerase
1.1.1.3: homoserine dehydrogenase
2.1.3.3: ornithine carbamoyltransferase
4.2.1.33: 3-isopropylmalate dehydratase

3.5.1.2: glutaminase
2.6.1.45: Transferases
2.5.1.54: DAHP synthase
4.2.1.10: dehydroquinase
2.7.1.71: shikimate kinase
2.7.1.39: homoserine kinase
4.2.3.1: threonine synthase
2.7.1.165:glycerate 2-kinase
4.1.3.27: anthranilate synthase
1.1.1.25:shikimate dehydrogenase
1.1.1.29:glycerate dehydrogenase
4.2.3.4:3-dehydroquinate synthase
4.3.2.1: argininosuccinate lyase
4.2.1.51: prephenate dehydratase

4.2.1.19: imidazoleglycerol-phosphate dehydratase
3.6.1.31: phosphoribosyl-ATP pyrophosphohydrolase
2.6.1.42: branched-chain amino acid aminotransferase
1.2.1.41: glutamate-5-semialdehyde dehydrogenase
3.5.1.18: succinyl-diaminopimelate desuccinylase
1.17.1.8: 4-hydroxy-tetrahydrodipicolinate reductase
1.2.1.38: N-acetyl-gamma-glutamyl-phosphate reductase
2.1.1.13: 5-methyltetrahydrofolate--homocysteine methyltransferase
5.3.1.16: phosphoribosylformimino-5-aminoimidazole carboxamide ribotide isomerase
5.4.2.12: 2,3-bisphosphoglycerate-independent phosphoglycerate mutase

2.6.1.51: serine---pyruvate transaminase
2.6.1.44: alanine-glyoxylate aminotransferase
2.6.1.17: succinyldiaminopimelate transaminase
2.3.1.31: homoserine O-acetyltransferase
1.1.1.85: 3-isopropylmalate dehydrogenase

Supplementary Fig. 10A

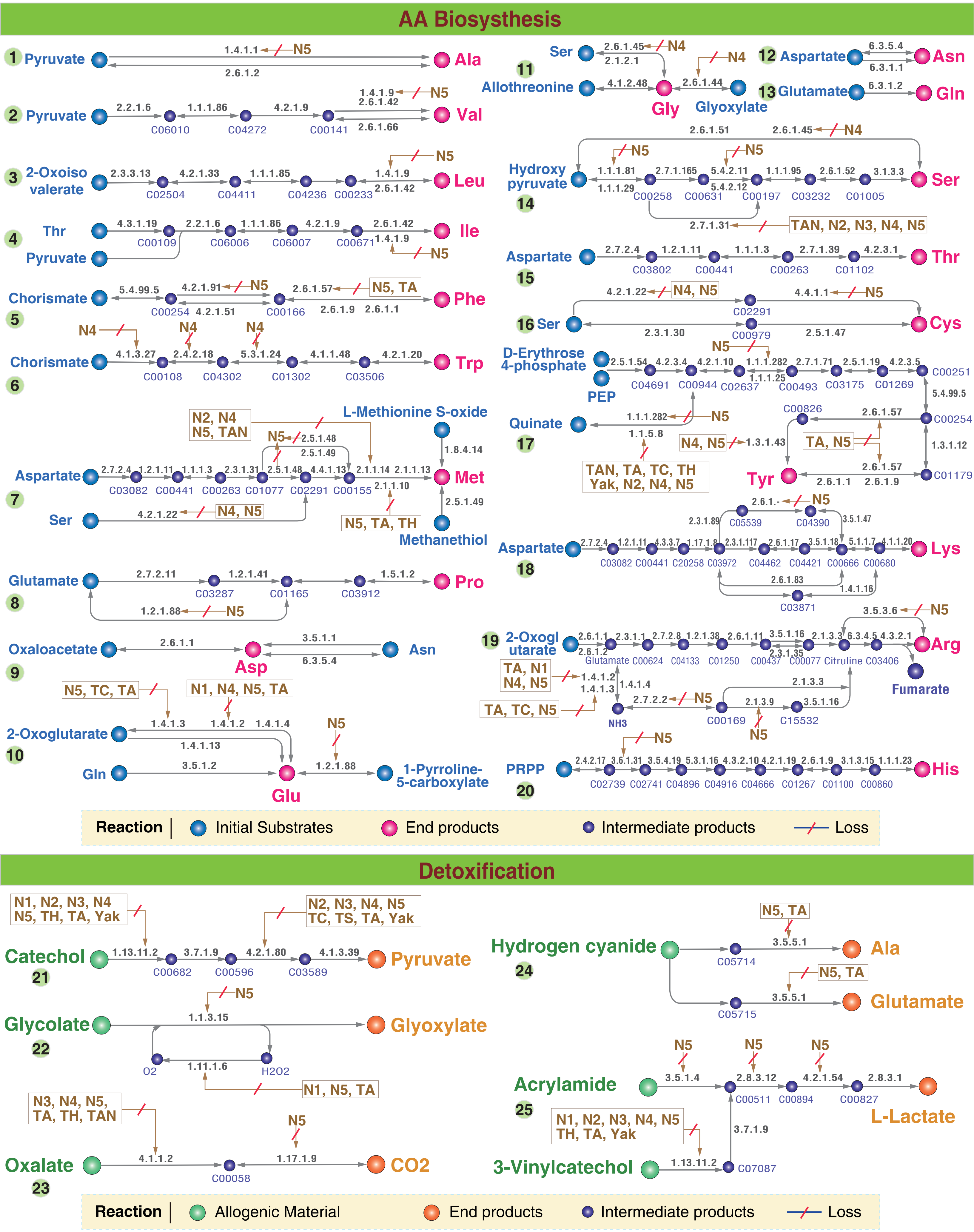

### Supplementary Fig. 10B

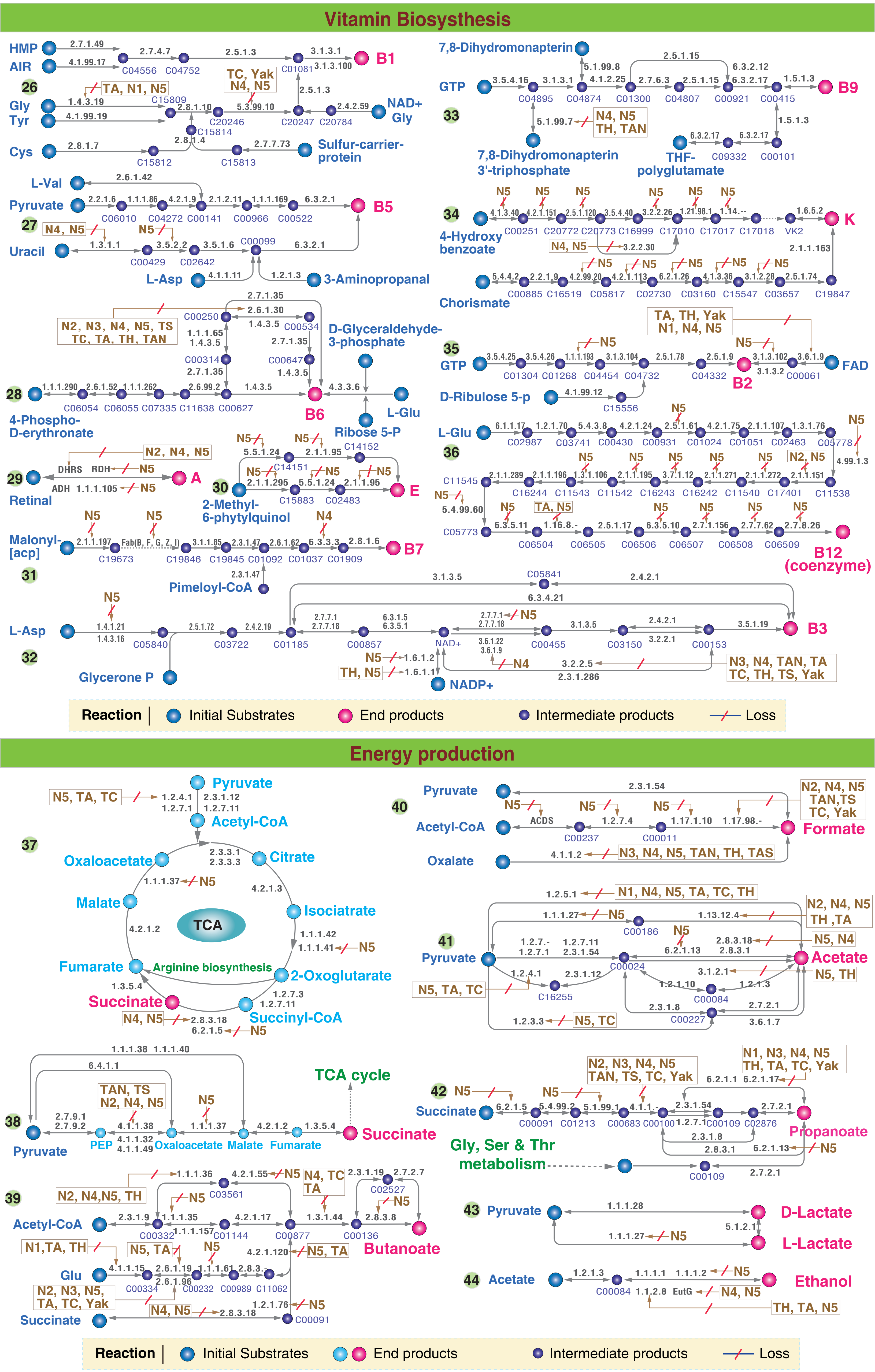

#### Supplementary Fig. 11

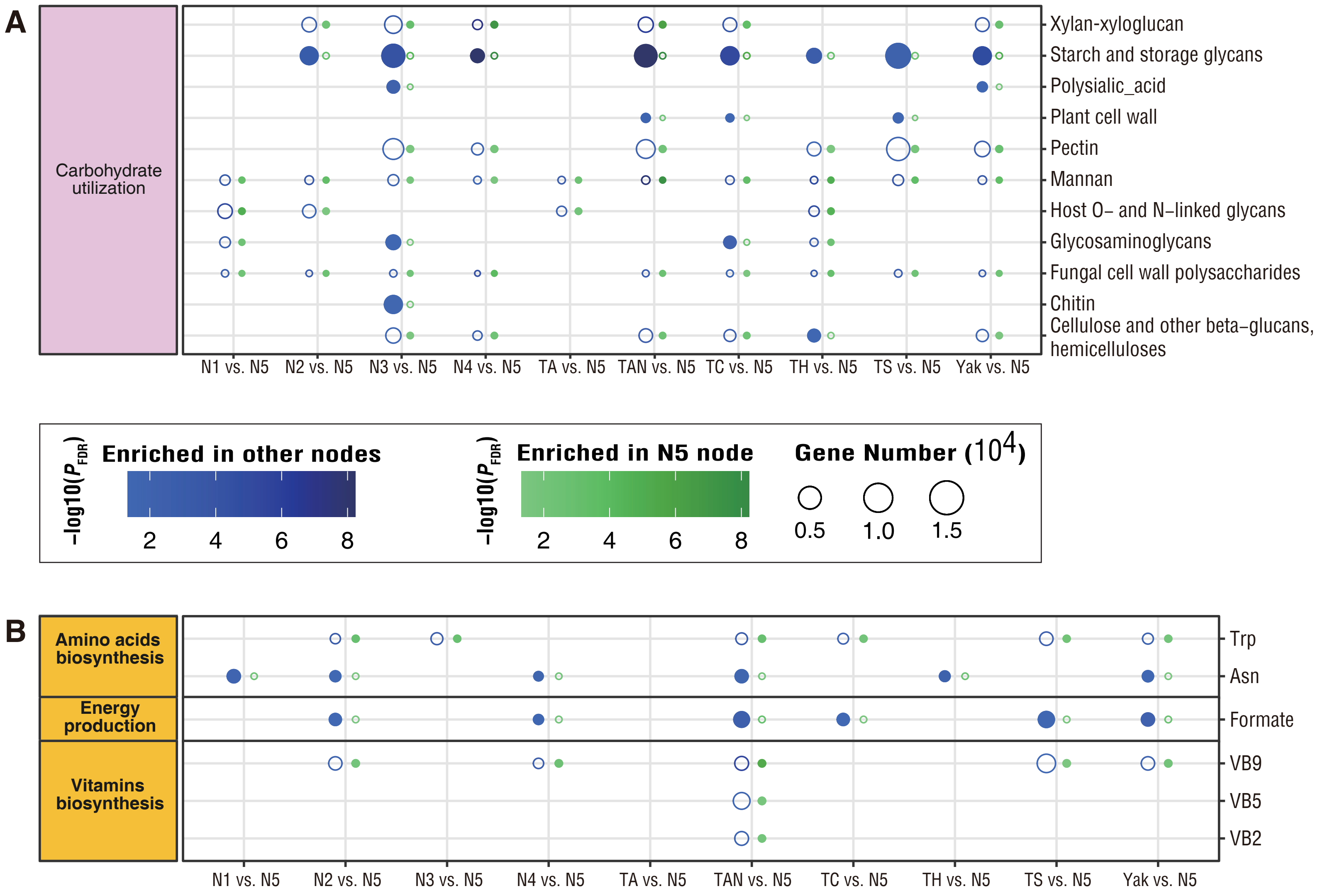

Supplementary Fig. 12

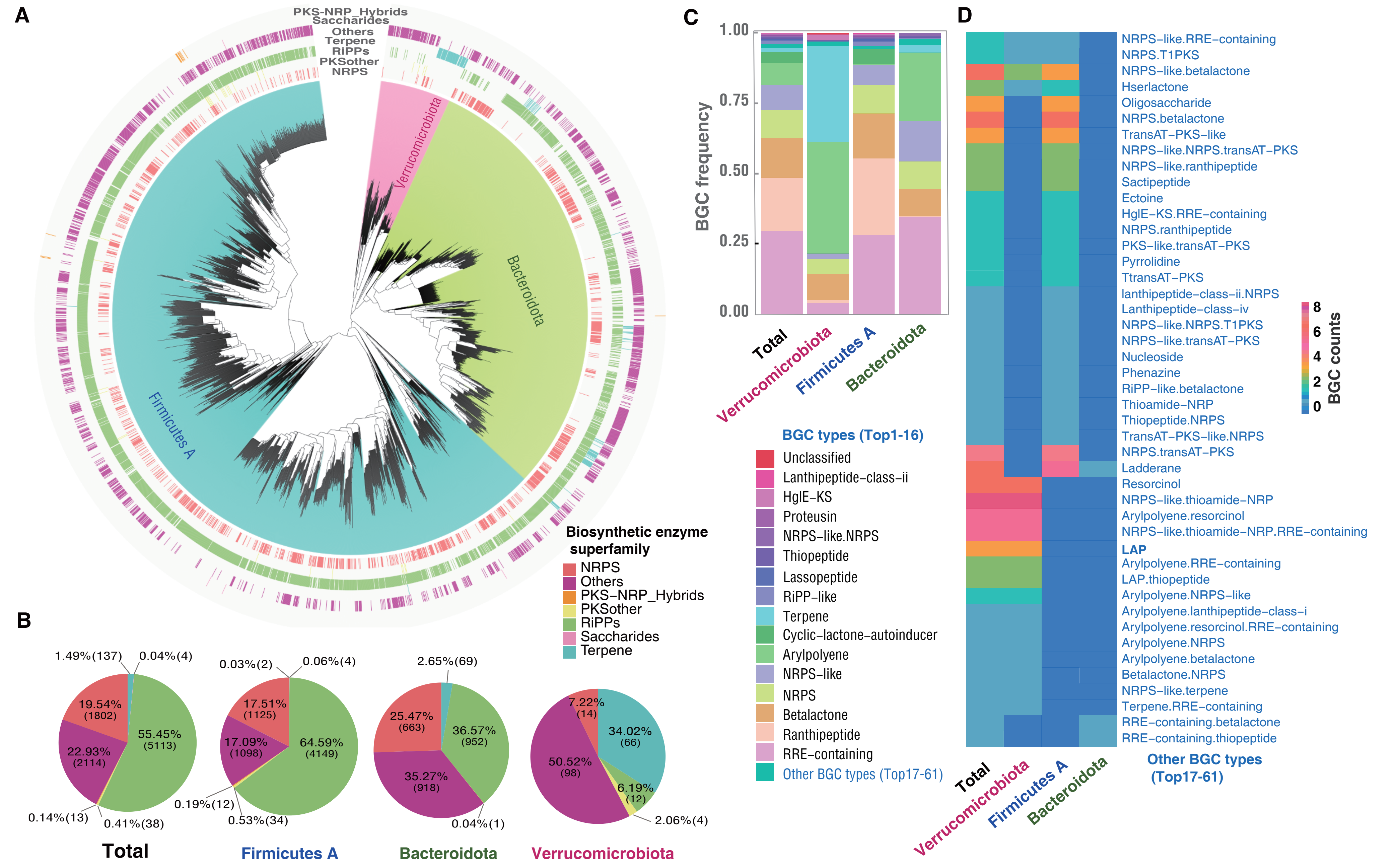
