## Supplementary information for "A Unified Catalog of 19,251 Non-human Reference Species Genomes Provides New Insights into the Mammalian Gut Microbiomes"

**Supplemental information**

This Supplementary Information file contains the following sections:

Material and Methods

Supplementary Texts

Supplementary Figure titles and summaries

Supplementary Table titles and summaries and

Reference.

**Materials and Methods**

### Sample collection

We collected fresh fecal samples from six native high-altitude mammals (including Tibetan Ass, Tibetan Horse, Tibetan Antelope, Tibetan Sheep and Tibetan Cattle) that have known phylogenetic relationships(*26-29*). Using the MGIEasy fecal sample collection kit (Cat. 10000035265, BGI, China), the inner parts of the fresh fecal samples of all 1,412 individuals were collected with a sterile spatula and then kept in the sterile sampling tubes while ensuring that our actions did not interfere with the animals' free lives in their natural habitat. The sampling time, animal species information, sample number and geographical location information were recorded (Fig. 1A). All fecal samples were stored in -80℃ freezers until transported to China National GeneBank (CNGB), Shenzhen on dry ice for further sequencing analysis. The detail information for the samples is shown in Supplementary Table 1.

### DNA extraction, sequencing and quality control

DNA extraction was performed using MagPure Stool DNA KF Kit B (Cat. MD5115, Magen) according to the standard kit protocol. DNA sequencing libraries were constructed by the MGIEasy Universal DNA Library Prep Set (Cat. 1000006986) and then sequenced on the DNBSEQ-T1 machine for 2 x 150 bp paired-end reads. A total of 33.52 Tb was produced from 1,412 samples and an average of 23.73 ± 7.22 Gb per sample clean reads were obtained after quality control by fastp v0.20.1(*44*) with default parameters. The Ass (GenBank assembly accession no. GCA_001305755.1), Tibetan cattle (GenBank assembly accession no. GCF_002263795.1), Tibetan horse (GenBank assembly accession no. GCA_002863925.1), Tibetan antelope (GenBank assembly accession no. GCA_000400835.1), Tibetan sheep (GenBank assembly accession no. GCA_002742125.1) and yak (GenBank assembly accession no. GCF_002263795.1) genomes were collected to build the host genomes index, and host contamination reads were removed by the Bowtie2 v2.3.5(*45*) alignments. Finally, an average of 23.55 ± 7.12 Gb high-quality reads per sample remained for subsequent analyses (Supplementary Table 2).

### Metagenomic assembly, binning and quality control

We assembled all the reads in each sample individually by the MEGAHITv1.2.9(*46*). For binning, a strategy that supports the co-abundance of contigs in multiple samples for better metagenomic binning has been proven(*47*), but has rarely been used in recent studies(*6-8, 32, 33*). In this study, we assessed the co-binning strategy according to tetranucleotide frequency, abundance correlation of contigs in multiple samples in MetaBAT2 v2.15(*48*) using the 79 Tibetan horse samples with different multiple sample size setting. We set the seven co-binning groups of 1, 4, 6, 8, 10, 15, and 20 samples (for convenience, they are referred to as s1 – s20), respectively, to test the influence of the different number of samples used in co-binning. Compared with the s1 group, the total number of bins, QS50 MAGs, and high-quality MAGs increased in the S4, S6, S8, S10, S15 and S20 groups (Supplementary Fig. 1A). In most of the samples, the number of QS50 MAGs per sample increased with the number of samples used in binning (Supplementary Fig. 1B). The QS50 MAGs in 7 groups were clustered into the species-level clusters by dRep V2.6.2(*49*) with the options: '--MASH_sketch 10000 --S_algorithm ANImf --P_ani 0.90 --S_ani 0.95 --cov_thresh 0.3' separately. More species were obtained when more samples were used in co-binning. Compared with s1, the total SGBs in the other groups were increased with the increase in sample numbers (Supplementary Fig. 1C). The overlaps of representative species genomes were calculated by dRep V2.6.2 in all the 7 groups. The result for estimation of co-binning strategy is shown in Supplementary texts. Considering both the consumption of computing resources and the SGBs yield, we selected 10 samples size for binning all the samples in this project. Subsequently, MetaBAT2 v2.15 (option '—min Contig 1500') was used to bin the assembled contigs into putative genomes within each sample. The resulting bins were referred to as metagenome-assembled genomes (MAGs). The completeness and contamination of MAGs were estimated using Check M v1.1.2(*50*) implemented in the 'lineage_wf' workflow. A total of 119,568 MAGs, called QS50 MAGs, passed the criteria defined by a previous study(*7*): >50% completeness, <10% contamination, and a quality score (completeness - 5x contamination, QS) > 50 (Supplementary Table 3). Next, the QS50 MAGs were further divided into 34,977 high-quality MAGs (completeness > 90% and contamination < 5%) and 84,591 medium-quality MAGs (others).

### Clustering MAGs into species-level genomes bins (SGBs)

All the 119,568 MAGs were clustered into strain-level genomes (ANI >= 99%) using dRep v2.6.2 with the options: '--MASH_sketch 10000 --S_algorithm ANImf --P_ani 0.95 --S_ani 0.99 --cov_thresh 0.3'. Representative genomes which have the maximum genome quality score: completeness - 5x contamination + 0.5log (N50) in each strain cluster were selected. The representative strain genomes were further clustered into the SGBs (ANI >= 95%) with the same options described above, except for '--P_ani 0.90 --S_ani 0.95'. All representative genomes in species clusters of each batch were merged and de-replicated using dRep V2.6.2 with the same options in the species-level step, and a non-human mammalian catalog consisting of 19,251 species-level reference genomes was reconstructed (Supplementary Table 4).

### Taxonomic annotations and phylogenetic analyses

Taxonomic annotation of 19,251 representative genomes of SGBs was performed with GTDB-Tk v1.4.1(*51*) and classified with the GTDB r95 database(*31*). The marker single copy proteins (122 for archaea, 120 for bacteria) were identified and the number of the marker genes in each SGBs was counted (Supplementary Table 4). In our SGBs, at least 38 marker genes were detected for taxonomic assignment (average 91±16). Phylogenetic tree of the non-human mammalian SGBs was generated using PhyloPhlAn v3.0.58(*52*) with the options "-d phylophlan --diversity high --fast --min_num_markers 80", and 18,607 SGBs with more than 80 markers were retained and visualized using GraPhlAn v1.1.4(*53*).

### Rarefaction curves

We constructed a rarefaction curve of the sample size and the number of SGB to assess whether the sample size is sufficient. The 19,251 SGBs’ curve was created by randomly re-sampling the pool of 1,412 samples 10 times with 100 sampling intervals, then all species and non-singleton species were plotted using the geom_smooth functions from the R ggplot2 package.

### Comparison with the public databases

To evaluate the novelty of our 19,251 SGB database, three public databases were used for the comparison.

1.	Rumen-Uncultured Genomes (RUG2) with 4,941 genomes(*32*). We used dRep V2.6.2 to cluster them into 2,177 species genomes.

2.	Genome Taxonomy Database (GTDB, r95) (*31*) with 31,910 species genomes.

3.	Genomes from Earth's Microbiomes catalog with 45,599 species genomes(*33*).

We calculated the genetic distance between our database and the other three databases using Mash v2.2.2(*54*) (options '-k 21 -s 10e4' for sketching) to evaluate overlaps between databases (Fig. 2C). To evaluate the reads mapping ratio of our samples, reads of each sample were re-sampled to 10M pairs, and then aligned to the four databases separately using Kraken2(*55*) with default parameters to calculate the mapping rates of each samples, which were visualized by the ‘ggplot2’ package in R (Fig. 2B).

### Computation of relative abundance for species genomes.

We used a similar strategy as described by Qin et. al.(*56*) to determine the presence or absence of specific SGBs and then calculated the relative abundance in each sample. In brief, to reduce the computational cost, reads of each sample were re-sampled to 10M paired-end reads. Next, the 19,251 species genomes were divided into 10 sub-databases, and reads were aligned to all sub-databases by bowtie2 (--end-to-end --sensitive -k 2), then unique alignments that met the following two criteria were retained: 1) both ends of reads aligned to the same contig; 2) identity of alignment was not lower than 0.95. To reduce the false-positive results, at least 100 paired-end reads were used to support the presence of given species in each sample and calculate their relative abundance using the same formula used in Qin et. al. (*56*).

### Diversity, PCoA and phylogenetic tree of microbial communities

Based on the relative abundance of SGBs in each sample, alpha diversity (both Richness and Shannon indices) was measured to estimate gut microbial diversity of the six hosts using the R vegan package. The Wilcoxon rank-sum test was used to evaluate the alpha diversity differences between host species (Fig. 3A). Beta diversity was calculated based on the Bray-Curtis dissimilarities between the 1,412 samples using the Vegan package in R, in which the ‘vegdist (method = “Bray”)’ was used, and then the principal coordinate analysis (PCoA) was performed with the ade4 package in R (Fig. 3C). We analyzed the paired Bray-Curtis dissimilarities among the samples from the levels of species, genera, families and classes (Fig. 3B). The Bray-Curtis dissimilarities between samples were hierarchically clustered by using the ‘hclust (method = “complete”)’ of the stats package in R, and the gut microbial community phylogenetic tree was constructed and compared with the host phylogenetic tree (Fig. 3D).

### Gene catalog building and functional annotations

We predicted protein-coding sequences (CDS) for 119,568 QS50 MAGs using Prodigal v2.6.3(*57*) with options '-c -m -p single'.Similar to the strategy for building genome catalogs, we constructed the gene sets of each host separately, and then combined 6 gene catalogs to obtain the non-human mammalian gut microbiome gene database. The function of the genes was annotated by using the Carbohydrate Active EnZyme (CAZy) Database and the Kyoto Encyclopedia of Genes and Genomes (KEGG) databases. The KEGG (V96) and NR (updated 2020_12_30) databases were selected for the protein sequence annotation with the blastp command in DIAMOND (v2.0.8.146) software(*58*). The CAZy database annotations were performed using the run_dbcan.py script with parameter "--tools all" in dbCAN2(*59*). All three state-of-the-art tools for CAZymes annotation were used: (ⅰ) HMMER search against the dbCAN HMM (hidden Markov model) database V9, (ⅱ) DIAMOND search against the CAZy pre-annotated CAZyme sequence database and (iii) Hotpep search against the conserved CAZyme short peptide (peptide pattern recognition) database. We combined all three outputs from these tools as the final CAZy annotation results.

We predicted biosynthetic gene clusters (BGCs) from our recalled SGBs using antiSMASH V6.0(*60*) based on profile hidden Markov models (pHMMs) from PFAM, TIGRFAMs or custom models. We then used BiG-SCAPE(*61*) to assign gene cluster families (GCFs) of the obtained BGCs. The results are presented in Supplementary Fig. 12 and Supplementary Table 13.

### Core microbiome of the six animal hosts and the ancestral microbiome reconstruction

The core microbiome was defined as those SGBs that were present in at least 50% of the samples in each animal host and can be identified using the ‘compute_core_microbiome.py’ script supplied in QIIME1(*62*).

To estimate the evolutionary dynamics in gut microbiomes over host evolutionary history, a presence/absence matrix of SGB from our identified core microbiome above was mapped against the host phylogeny and gain-and-loss patterns of SGBs along the host phylogeny were determined using the asymmetrical Wagner parsimony approach in Count v.10.04(*36*) with a gain penalty of 1.0. The reliability of similar ancestral microbiome reconstruction has been confirmed by a previous study of gut microbiomes(*63*).

### Co-phylogeny analysis between host species and gut microbial individual species/strain

The prevalence of specifically gained SGBs and the shared SGBs in the six hosts were counted and the significance of the difference was calculated by the Wilcoxon rank sum test (Fig. 4B). Considering the evolutionary trait, the bacterial genera including the specifically gained SGBs from at least four hosts were chosen to perform the co-phylogeny analysis. In total, phylogenetic trees (Supplementary Fig. 7) of eight genera from three representative phyla meeting those conditions were generated using PhyloPhlAn v3.0.58(*52*) (parameters: "-d phylophlan --diversity high --accurate --min_num_markers 80") and visualized by using iTol v6.4.3(*64*). The frequent swap events of SGBs across the six hosts were further visualized by the R ‘circlize’ package (Fig. 4C). To view the host restriction for the bacterial strains, the SGBs supporting MAGs that were assembled from at least four hosts were chosen, and only two SGBs (SGB1640 and SGB798) fitted this condition. The Phylogenetic trees of these two SGBs were built using the method described above (Supplementary Fig.8).

### Functional analyses

Using the annotated results from KEGG and CAZy databases, we first investigated the common functional traits of AFBs including carbohydrate utilization, energy production& key precursors, amino acid biosynthesis, and vitamin biosynthesis (Fig. 5A, Supplementary Fig. 9; Supplementary Table 10). The abilities to produce specific metabolites by the AFBs were determined by the presence of at least one complete reaction pathway using the combination of the results of six AFBs (Supplementary Fig. 10, Supplementary Table 10).

Next, the functional enrichment analysis at the pathway levels of lineage-specific gained SGBs among different evolutionary nodes (N1-N5) (Fig.5B, Fig.6, Supplementary Fig. 11 and Supplementary Table 11 and 12) was based on the Fisher's exact test (one-sided way) in R. Based on the functionally annotated results from KEGG and CAZy, the number of genes annotated to different KEGG Orthologs (KOs) in each SGB was counted. To determine functional enrichments at the KEGG pathway level of the specifically gained SGBs at each evolutionary node, we counted the number of genes annotated to each pathway. The function matrix (c(A, B, C, D), nrow=2) in R was used to represent the matrix of gene numbers between any two compared nodes in a particular pathway. Where A-D represents the gene numbers present or absent in a given pathway of two compared nodes, respectively. The fisher.test() function in R was used to compute the significance of the enrichment with parameters (alternative="less" or "greater"). For each given pathway, the model ("less" or "greater") with a lower probability value in the Fisher-exact test was chosen to determine the significance and enrichment direction.

### Statistical analysis

Statistical significance was verified through Fisher’s test or Wilcoxon rank sum test, and multiple hypothesis testing correction was performed using the false discovery rate (FDR) methods. Adjusted-P values lower than 0.05 were considered to be statistically significant. All other computational and statistical analyses were performed with the open-source software tools that have been cited in the methods.

**Supplementary Texts**

### Assembly statistics

A series of read quality control and assembly pipelines were used (Methods), and finally we acquired an average of 2.73 million contigs (±0.84) with a length of more than 200 bp for each sample (N50: 833±292) (Supplementary Table 2).

### Estimation of co-binning strategy

Compared to the single-sample binning result, the total binning number, the QS50 MAGs, and the average MAGs per sample were upraised over 147.17% in the minimum samples size setting (S4) (Supplementary Fig. 1A and 1B). Furthermore, the rate of all the indices increased as the sample size increased (Supplementary Fig. 1A and 1B). After the MAGs were clustered according to a species threshold of average nucleotide identity (ANI) of ≥ 95% (Methods), more species-level genome bins (SGBs) (at least 125.64% increasing rate in S4) were obtained by this method (Supplementary Fig. 1C-D). Considering both the availability of computing resources and the yield of SGBs, we selected and performed the co-binning of contigs assembled from every ten samples in this project because of 174.93% increase in the bins and 153.53% increase for the SGBs in the test samples.

### Quality of MAGs

The high-quality genome size of our MAGs was from 0.47M~6.14M and the GC content (%) ranged from 23.75%~72.59% (Supplementary Table 3). There were 23 near complete MAGs having 100% completeness and zero contamination, suggesting that our assembled genomes were highly reliable for further analyses.

### Non-species taxonomic assignments of 19,251 SGBs

There were 14,039 (73.64%) SGBs classified to 583 known genera, 18,751 (98.34%) SGBs to 180 known families, and 19,012 (99.71%) SGBs to 90 known orders and all the SGBs had clear classification information at the 42 known classes and 25 known phyla. A total of 1,734 SGBs were identified as novel genera and 27 SGBs belonged to novel families.

### Analysis of Alpha and beta diversity

Based on the SGB profiling, we found that the species richness of the gut microbiome differed significantly even between the two phylogenetically closest species, Tibetan cattle and Yak (Fig. 3A) and the same trend was observed for the Shannon index except for the comparison between yak and cattle where no significant difference was observed (Fig. 3A) suggesting a complex interaction between host genetics and gut microbiota diversity. Still, a pairwise comparison demonstrated that the Bray-Curtis dissimilarity increased as the phylogenetic differences between the hosts increased pointing to an effect of host genetics(Fig. 3B), a notion also supported by principal coordinates analysis (PCoA) of beta diversity (Fig. 3C and Supplementary Fig. 4). Further analysis of the phylogenetic tree of microbial communities based on Bray-Curtis dissimilarity using all the SGBs and core SGBs (defined as those at least occurred in 50% samples per host) showed the same [topological structure](javascript:;) compared with host phylogeny (Fig. 3D and Supplementary Fig. 5).

### Functional divergences of HSG SGBs

Two paired comparisons of N1 (the common ancestor between Tibetan horse and Tibetan Ass) vs. N4 (the common ancestor between Bovinae and Caprinae) and N2 (the common ancestor between Tibetan sheep and Tibetan antelope) vs. N3 (the common ancestor between Tibetan cattle and Yak) showed consistent differentiation of multiple metabolic pathways, such as the ability to metabolize multiple polysaccharides, starch and storage glycans, mannan, xylan, amino acids, vitamins and co-factors, which is independent of host divergence time and may be related to specific environmental stress. However, the mechanism remains unclear (Fig. 6A). Moreover, significant functional changes of HSG SGBs occurred uniquely to the divergence between nodes N2 and N3, including the hydrolysis of plant cell wall polysaccharides, pyruvate metabolism, lysine biosynthesis, cysteine and methionine metabolism. Similarly, unique to the comparison between nodes N1 and N4, we also observed significant differences in a number of enriched pathways: phenylalanine metabolism, histidine metabolism, biosynthesis of ubiquinone and other terpenoid−quinones.

**Supplementary Figure titles and summaries**

### Supplementary Fig. 1. The effective estimation of co-binned MAGs and SGBs using all 79 Tibetan horse samples.

(**A**) The effect of co-binning sample size setting on the obtained MAG number. (**B**) The effect of co-binning sample size setting on the QS50 MAGs per sample. The node of the same sample is strung by the grey line. (**C**) Estimation of effective SGBs improvement in different sample sizes. (**D**) Overlapping of effective SGBs obtained from different sample size settings.

### Supplementary Fig. 2. Quality assessment of 19,251 SGBs.

(**A**)The completeness and contamination of 19,251 SGBs. If the SGB satisfied the criteria (>90% completeness and <5% contamination), the SGB is identified as high quality and marked by the right dots. (**B**) Genome sizes and (**C**) N50 statistics of SGBs. The blue box stands for the medium MAGs and the red box stands for the high quality. (**D**)The distribution of MAG numbers supporting SGBs.

### Supplementary Fig. 3. Taxonomic statistics of 19,251 SGBs.

(**A**) The taxonomic classification of the bacteria in the 19,251 SGBs. The species classification is performed based on the GTDB database. Only the top five most frequently observed taxa are shown in the figure and the remaining lineages are shown as “others”. The white pillars show the proportion of novel SGBs in each rank level. (**B**) The taxonomic classification of the archaea in the 19,251 SGBs. The species classification is performed based on the GTDB database. Only the top three most frequently observed taxa are shown in the figure and the remaining lineages were shown as “others”. The white pillars show the novel SGBs proportion in each rank level.

### Supplementary Fig. 4. The PCoA analysis in three paired hosts.

(**A**) TA and TH; (**B**) TAN and TS; (**C**) TC and Yak.

### Supplementary Fig. 5. Core microbial community tree based on Bray-Curtis dissimilarity.

Microbial community tree constructed using the Core 50 microbiota by the Bray-Curtis dissimilarity shows the phylosymbiotic pattern with the host phylogeny. The right is the phylogenetic tree of the six host animal species. The Core 50 SGBs numbers and the sample numbers in each host are shown in the brackets.

### Supplementary Fig. 6. The phylogenetic tree of core 50 SGBs belonging to three representative bacterial phyla and the gained SGBs in different host evolutionary nodes. Linked to Figure 4A.

The phylogenetic tree of core 50 SGBs belonging to three representative bacterial phyla was constructed using the Maximum-likelihood method suppled in PhyloPhlAn software (See Supplementary Methods). The color of the branch stands for the classification of the genus. The genera signed by red stars are selected to perform the next co-speciation analysis. The core microbial forming profiling over host evolutionary history is shown based on the phylogenetic tree of SGBs from the Firmicute A (**A**), Bacteroidota (**B**) and Verrucomicrobiota (**C**) phyla. The AFBs (SGB_2397, SGB_5789, SGB_13840, SGB_18780, SGB_18778, and SGB_18783) are showed in the phylogenetic tree.

### Supplementary Fig. 7. Maximum-likelihood phylogeny of the representative eight genera from tree phyla indicate host phylogenetic restriction (Linked to Figure 4C).

Maximum-likelihood phylogeny of the selected 8 core genera from three representative bacterial phyla. (**A**) *CGA-110*, (**B**) *Ruminococcus*, (**C**) *Agathobacter* and (**D**) *Acetatifactor* from Firmicutes A. (**E**) *RC9* and (**F**) *Alitipes* from the Bacteroidota. (**G**) *Akkermansia* and (**H**) *UBA1067* from the Verrucomicrobiota. The outgroup included five SGBs from Cyanobacteria phylum (an ancient bacterial phylum that is the closest to the Firmicutes) covering the most broadly conserved microbial gene markers used to build phylogenetic trees. Light and dark blue dots in the branch of each tree only show the bootstrap values higher than 50% and 70% supporting topological structures, respectively.

### Supplementary Fig. 8. Host phylogenetic restriction strain tree.

Maximum-likelihood phylogenetic trees of the SGB stains assembled from more than four hosts. (**A**) SGB1640 and (**B**) SGB798. Light and dark blue dots in the branch of each tree only show the bootstrap values higher than 50% and 70% supporting topological structures, respectively.

### Supplementary Fig. 9. Detailed reaction steps for amino acid (AA) biosynthesis of six founder bacteria linked to Figure 5A.

The reaction steps for biosynthesis of the 20 AA are shown. The enzyme code is marked on the reaction chain, and the colors stand for the enzymes that can be found in the corresponding AFB. Except for histidine, every enzyme step of the other AAs is found in at least one AFB. See details in Supplementary Table 10.

### Supplementary Fig. 10. Metabolic reactions of 6 founders at N5 and gained bacteria at other nodes linked to Fig. 5B.

Based on the complete metabolic pathways, the presence of enzymes of the N5 and gained bacteria at other nodes are shown. The red slash on the arrow line implies the enzyme was not found in the group. (**A**) Vitamin biosynthesis and energy production. (**B**) Amino acid biosynthesis and detoxification. See details in Supplementary Table 10.

### Supplementary Fig. 11. Comparison of enrichment between founder node and other nodes.

Using the N5 founder node as the reference, the enrichments of other nodes are shown. The size of the circle represents the number of genes associated with substrate utilization capacities or the metabolic pathways. The solid circle represents the enriched node. The color scale from light to dark corresponds to the negative logarithm of FDR-adjusted P values from low to high. (**A**) Carbohydrate utilization. (**B**) Amino acids biosynthesis, Energy production and Vitamin biosynthesis. See details in Supplementary Table 11.

### Supplementary Fig. 12. Predicted biosynthetic genes from three core bacterial phyla Firmicute A, Bacteroidota, and Verrucomicrobiota.

From 3888 (69.34%) of 5607 SGBs of the three core bacterial phyla (the SGB tree is built according to the method described in Extended Data Fig. 6), we identified 9,221 BGCs (including 130,098 intact CDSs) belonging to seven biosynthetic enzyme superfamilies, which were distributed distinctly into the three bacterial phyla (**A-B**). The 9,218 of these BGCs were assigned into 60 known BGC types and showed different dominances in the three bacterial phyla (**C-D**). See details in Supplementary Table 13.

**Supplementary Table titles and summaries**

### Supplementary Table 1. Information on samples included in the study.

### Supplementary Table 2. Data summary of the samples.

### Supplementary Table 3. Information of the assembled MAGs in the study.

### Supplementary Table 4. Summary of SGBs in the study.

### Supplementary Table 5. Profiling of 19,251 SGBs from the 1,412 samples.

### Supplementary Table 6. Data summary of the genome and gene catalog of the gut microbiomes.

(A) Summary of the 19,251 assembled genome catalog. (B) Summary of the gene catalog.

### Supplementary Table 7. The expanding diversity of species at the phylum level.

### Supplementary Table 8. SGBs information of the eight core bacterial genera used for the co-phylogeny analysis.

### Supplementary Table 9. Strain information of two bacterial species used for the co-phylogeny analysis.

### Supplementary Table 10. Functional information of AFBs and gained SGBs linked to Figs 5, Fig. 6 and Supplementary Figs. 9-11.

(A) KEGG annotated results and (B) CAZy annotated results.

### Supplementary Table 11. Enrichment analysis of non-AFB nodes compared with AFB node linked to Figs 5B, Supplementary Fig. 11 and Table S10.

### Supplementary Table 12. Functional enrichment of gained SGBs in different evolutionary nodes linked to Fig.6 and Table S10.

(A) Enrichment of KEGG pathways and (B) Carbohydrate utilization.

### Supplementary Table 13. Basic summary of 9,221 predicted biosynthetic gene clusters (BGCs) linked to Supplementary Fig. 12.

**References**

1.	Z. Zhang *et al.*, Convergent Evolution of Rumen Microbiomes in High-Altitude Mammals. *Current Biology* **26**, 1873-1879 (2016).

2.	P. Rosshart Stephan *et al.*, Laboratory mice born to wild mice have natural microbiota and model human immune responses. *Science* **365**, eaaw4361 (2019).

3.	C. Campbell *et al.*, Bacterial metabolism of bile acids promotes generation of peripheral regulatory T cells. *Nature* **581**, 475-479 (2020).

4.	S. K. Gill, M. Rossi, B. Bajka, K. Whelan, Dietary fibre in gastrointestinal health and disease. *Nature Reviews Gastroenterology & Hepatology* **18**, 101-116 (2021).

5.	P. Kundu, E. Blacher, E. Elinav, S. Pettersson, Our Gut Microbiome: The Evolving Inner Self. *Cell* **171**, 1481-1493 (2017).

6.	E. Pasolli *et al.*, Extensive Unexplored Human Microbiome Diversity Revealed by Over 150,000 Genomes from Metagenomes Spanning Age, Geography, and Lifestyle. *Cell* **176**, 649-662.e620 (2019).

7.	A. Almeida *et al.*, A unified catalog of 204,938 reference genomes from the human gut microbiome. *Nature Biotechnology* **39**, 105-114 (2021).

8.	D. Levin *et al.*, Diversity and functional landscapes in the microbiota of animals in the wild. *Science* **372**, eabb5352 (2021).

9.	J. Qiu, China: The third pole. *Nature* **454**, 393-396 (2008).

10.	Z. Zhou, T. Deng, The Tibetan Plateau is a natural laboratory for studying organic evolution and environmental change. *Science China Earth Sciences* **63**, 169-171 (2020).

11.	T. Deng, F. Wu, Z. Zhou, T. Su, Tibetan Plateau: An evolutionary junction for the history of modern biodiversity. *Science China Earth Sciences* **63**, 172-187 (2020).

12.	T. Deng *et al.*, Out of Tibet: Pliocene Woolly Rhino Suggests High-Plateau Origin of Ice Age Megaherbivores. *Science* **333**, 1285-1288 (2011).

13.	L. Cortes-Ortiz, K. R. Amato, Host genetics influence the gut microbiome. *Science* **373**, 159-160 (2021).

14.	A. H. Moeller, T. A. Suzuki, M. Phifer-Rixey, M. W. Nachman, Transmission modes of the mammalian gut microbiota. *Science* **362**, 453-457 (2018).

15.	L. Grieneisen *et al.*, Gut microbiome heritability is nearly universal but environmentally contingent. *Science* **373**, 181-186 (2021).

16.	P. Ferretti *et al.*, Mother-to-Infant Microbial Transmission from Different Body Sites Shapes the Developing Infant Gut Microbiome. *Cell Host & Microbe* **24**, 133-145.e135 (2018).

17.	R. M. Brucker, S. R. Bordenstein, The roles of host evolutionary relationships (genus: Nasonia) and development in structuring microbial communities. *Evolution* **66**, 349-362 (2012).

18.	M. Groussin, F. Mazel, E. J. Alm, Co-evolution and Co-speciation of Host-Gut Bacteria Systems. *Cell Host & Microbe* **28**, 12-22 (2020).

19.	F. Mazel *et al.*, Is Host Filtering the Main Driver of Phylosymbiosis across the Tree of Life? *mSystems* **3**, e00097-00018 (2018).

20.	M. Groussin *et al.*, Unraveling the processes shaping mammalian gut microbiomes over evolutionary time. *Nature Communications* **8**, 14319 (2017).

21.	H. Moeller Andrew *et al.*, Cospeciation of gut microbiota with hominids. *Science* **353**, 380-382 (2016).

22.	A. Gaulke Christopher *et al.*, Ecophylogenetics Clarifies the Evolutionary Association between Mammals and Their Gut Microbiota. *mBio* **9**, e01348-01318 (2018).

23.	N. D. Youngblut *et al.*, Host diet and evolutionary history explain different aspects of gut microbiome diversity among vertebrate clades. *Nature Communications* **10**, 2200 (2019).

24.	J. R. Brown, C. J. Douady, M. J. Italia, W. E. Marshall, M. J. Stanhope, Universal trees based on large combined protein sequence data sets. *Nat Genet* **28**, 281-285 (2001).

25.	F. Delsuc, H. Brinkmann, H. Philippe, Phylogenomics and the reconstruction of the tree of life. *Nature Reviews Genetics* **6**, 361-375 (2005).

26.	H. Jonsson *et al.*, Speciation with gene flow in equids despite extensive chromosomal plasticity. *Proc Natl Acad Sci U S A* **111**, 18655-18660 (2014).

27.	Y. Jiang *et al.*, The sheep genome illuminates biology of the rumen and lipid metabolism. *Science* **344**, 1168-1173 (2014).

28.	L. Chen *et al.*, Large-scale ruminant genome sequencing provides insights into their evolution and distinct traits. *Science* **364**, eaav6202 (2019).

29.	A. M. Humphreys, T. G. Barraclough, The evolutionary reality of higher taxa in mammals. *Proceedings. Biological sciences* **281**, 20132750 (2014).

30.	D. H. Parks *et al.*, Recovery of nearly 8,000 metagenome-assembled genomes substantially expands the tree of life. *Nature Microbiology* **2**, 1533-1542 (2017).

31.	D. H. Parks *et al.*, GTDB: an ongoing census of bacterial and archaeal diversity through a phylogenetically consistent, rank normalized and complete genome-based taxonomy. *Nucleic acids research* **50**, D785-D794 (2022).

32.	R. D. Stewart *et al.*, Compendium of 4,941 rumen metagenome-assembled genomes for rumen microbiome biology and enzyme discovery. *Nature Biotechnology* **37**, 953-961 (2019).

33.	S. Nayfach *et al.*, A genomic catalog of Earth’s microbiomes. *Nature Biotechnology* **39**, 499-509 (2021).

34.	L. Glendinning, B. Genç, R. J. Wallace, M. Watson, Metagenomic analysis of the cow, sheep, reindeer and red deer rumen. *Scientific Reports* **11**, 1990 (2021).

35.	F. Xie *et al.*, An integrated gene catalog and over 10,000 metagenome-assembled genomes from the gastrointestinal microbiome of ruminants. *Microbiome* **9**, 137 (2021).

36.	M. Csurös, Count: evolutionary analysis of phylogenetic profiles with parsimony and likelihood. *Bioinformatics (Oxford, England)* **26**, 1910-1912 (2010).

37.	K. Kwong Waldan *et al.*, Dynamic microbiome evolution in social bees. *Science Advances* **3**, e1600513 (2017).

38.	D. M. de Vienne *et al.*, Cospeciation vs host-shift speciation: methods for testing, evidence from natural associations and relation to coevolution. *New Phytologist* **198**, 347-385 (2013).

39.	Q. Qiu *et al.*, The yak genome and adaptation to life at high altitude. *Nat Genet* **44**, 946-949 (2012).

40.	R.-L. Ge *et al.*, Draft genome sequence of the Tibetan antelope. *Nat Commun* **4**, 1858 (2013).

41.	L. Yu, Y. Chen, W. Wang, Z. Xiao, Y. Hong, Multi-Vitamin B Supplementation ReversesHypoxia-Induced Tau Hyperphosphorylation and Improves Memory Function in Adult Mice. *Journal of Alzheimer's Disease* **54**, 297-306 (2016).

42.	Y. P. Wang *et al.*, Riboflavin supplementation improves energy metabolism in mice exposed to acute hypoxia. *Physiol Res* **63**, 341-350 (2014).

43.	S. C. Nunes *et al.*, Cysteine boosters the evolutionary adaptation to CoCl2 mimicked hypoxia conditions, favouring carboplatin resistance in ovarian cancer. *BMC Evolutionary Biology* **18**, 97 (2018).

44.	S. Chen, Y. Zhou, Y. Chen, J. Gu, fastp: an ultra-fast all-in-one FASTQ preprocessor. *Bioinformatics (Oxford, England)* **34**, i884-i890 (2018).

45.	B. Langmead, S. L. Salzberg, Fast gapped-read alignment with Bowtie 2. *Nature Methods* **9**, 357-359 (2012).

46.	D. Li, C.-M. Liu, R. Luo, K. Sadakane, T.-W. Lam, MEGAHIT: an ultra-fast single-node solution for large and complex metagenomics assembly via succinct de Bruijn graph. *Bioinformatics (Oxford, England)* **31**, 1674-1676 (2015).

47.	J. N. Nissen *et al.*, Improved metagenome binning and assembly using deep variational autoencoders. *Nature Biotechnology* **39**, 555-560 (2021).

48.	D. D. Kang *et al.*, MetaBAT 2: an adaptive binning algorithm for robust and efficient genome reconstruction from metagenome assemblies. *PeerJ* **7**, e7359-e7359 (2019).

49.	M. R. Olm, C. T. Brown, B. Brooks, J. F. Banfield, dRep: a tool for fast and accurate genomic comparisons that enables improved genome recovery from metagenomes through de-replication. *The ISME Journal* **11**, 2864-2868 (2017).

50.	D. H. Parks, M. Imelfort, C. T. Skennerton, P. Hugenholtz, G. W. Tyson, CheckM: assessing the quality of microbial genomes recovered from isolates, single cells, and metagenomes. *Genome Res* **25**, 1043-1055 (2015).

51.	P.-A. Chaumeil, A. J. Mussig, P. Hugenholtz, D. H. Parks, GTDB-Tk: a toolkit to classify genomes with the Genome Taxonomy Database. *Bioinformatics (Oxford, England)* **36**, 1925-1927 (2020).

52.	N. Segata, D. Börnigen, X. C. Morgan, C. Huttenhower, PhyloPhlAn is a new method for improved phylogenetic and taxonomic placement of microbes. *Nature Communications* **4**, 2304 (2013).

53.	F. Asnicar, G. Weingart, T. L. Tickle, C. Huttenhower, N. Segata, Compact graphical representation of phylogenetic data and metadata with GraPhlAn. *PeerJ* **3**, e1029 (2015).

54.	B. D. Ondov *et al.*, Mash Screen: high-throughput sequence containment estimation for genome discovery. *Genome Biol* **20**, 232-232 (2019).

55.	D. E. Wood, J. Lu, B. Langmead, Improved metagenomic analysis with Kraken 2. *Genome Biol* **20**, 257-257 (2019).

56.	J. Qin *et al.*, A metagenome-wide association study of gut microbiota in type 2 diabetes. *Nature* **490**, 55-60 (2012).

57.	D. Hyatt *et al.*, Prodigal: prokaryotic gene recognition and translation initiation site identification. *BMC Bioinformatics* **11**, 119 (2010).

58.	B. Buchfink, C. Xie, D. H. Huson, Fast and sensitive protein alignment using DIAMOND. *Nat Meth* **12**, 59-60 (2015).

59.	H. Zhang *et al.*, dbCAN2: a meta server for automated carbohydrate-active enzyme annotation. *Nucleic Acids Research* **46**, W95-W101 (2018).

60.	K. Blin *et al.*, antiSMASH 6.0: improving cluster detection and comparison capabilities. *Nucleic Acids Research* **49**, W29-W35 (2021).

61.	J. C. Navarro-Muñoz *et al.*, A computational framework to explore large-scale biosynthetic diversity. *Nature chemical biology* **16**, 60-68 (2020).

62.	J. G. Caporaso *et al.*, QIIME allows analysis of high-throughput community sequencing data. *Nature Methods* **7**, 335-336 (2010).

63.	W. K. Kwong *et al.*, Dynamic microbiome evolution in social bees. *Science Advances* **3**, e1600513 (2017).

64.	I. Letunic, P. Bork, Interactive Tree Of Life (iTOL) v5: an online tool for phylogenetic tree display and annotation. *Nucleic acids research* **49**, W293-W296 (2021).
